# Molecular Insights into Cell Wall Architecture and Xylan-Bound Cellulose Fibrils in the Wheat Straw

**DOI:** 10.1101/2025.03.11.642666

**Authors:** Yucheng Hu, Shixu Yu, Zhe Ling, Peng Xiao, Yutong Zhu, Guohua Miao, Yuan He, Haichao Li, Sheng Chen, Tingting You, Tuo Wang, Feng Xu

**Affiliations:** State Key Laboratory of Efficient Production of Forest Resources, Beijing Key Laboratory of Lignocellulosic Chemistry, Beijing Forestry University, Beijing 100083, China; Jiangsu Co-Innovation Center of Efficient Processing and Utilization of Forest Resources, College of Chemical Engineering, Nanjing Forestry University, Nanjing 210037, China; Department of Chemistry, Michigan State University, East Lansing, MI 48824, USA

## Abstract

The plant cell wall, a highly abundant renewable resource, remains insufficiently understood in terms of its structure, particularly the architecture of cellulose microfibrils and their interactions with hemicellulose and lignin, which impede the development of efficient downstream conversion. Here we present an alternative model of xylan-bound cellulose microfibril architecture in the cell wall of intact wheat straw, using solid-state nuclear magnetic resonance (ssNMR) coupled with X-ray scattering techniques. We show that the spectroscopic and scattering data can be fit to a simple elementary microfibril consisting of 18 glucan chains, with additional contributions from 0 to 4 tightly bound flat-ribbon xylan chains across the cellulose fibril axis. These xylan-bound cellulose microfibrils are found to serve as the secondary interaction site with lignin, following the non-flat xylan domains. The loose packing of these xylan-bound cellulose fibrils results in a heterogenous hydration landscape within the cell wall, allowing water penetration while maintaining structural integrity through numerous physical contacts between the polymers. This study provides a potential framework to reconcile the mounting biochemical evidence supporting the 18-chain elementary cellulose microfibril model and cellulose synthase complex with the oversized observations from characterization techniques, which report averaged structures influenced by microfibril coalescence and hemicellulose binding. The structural insights also offer valuable information regarding recalcitrance and the potential applications of biomass in biorefinery operations.

## Introduction

Lignocellulosic materials from grasses, such as maize, wheat straw, biomass sorghum, and rice straw, are among the major feedstocks used for the production of biofuels, biopolymers, and bio-based chemicals^1–3^. he grass cell wall features a supramolecular network in both primary and secondary walls, with cellulose, hemicellulose, and lignin heterogeneously distributed, providing structural support while protecting the plant from herbivory and pathogens. However, this complex architecture also makes the cell wall inherently recalcitrant, significantly hindering the efficient depolymerization of plant cell walls into monomers^4^. Understanding the molecular architecture of grass cell walls is crucial for advancing biomass process technology and developing genetically engineered plants^5^.

The assembly and structure of the plant cell wall is regulated by numerous molecular interactions among cellulose, hemicellulose, and lignin. Cellulose consists of β-1,4 glucan chains with the conformation of flat two-fold screw ribbons. While the number of glucan chains within an elementary microfibril has been debated, it is mostly likely to contain 18, 24, or 36 chains^6–8^. Mounting biochemical, structural, and modeling data from studies on the cellulose synthase complex and microfibrils synthesized in vitro and found in the cell wall converge to support the 18-chain model^6,9–12^. Coalescence of multiple elementary microfibrils commonly occurs, influencing the mechanical properties of the cell wall^13^, and may contribute to the discrepancy in chain number counts observed in characterization techniques, alongside the deposition of structured xylan molecules on the fibril surface^14^. Xylan, a hemicellulose widely found in grass cell walls, has a backbone made of linear β-1,4-linked xylose residues, which are substituted with α-1,3-linked arabinose (Ara), α-1,2-linked glucuronic acid (GlcA), and often acetylated at the O-2 and O-3 positions^15^. Xylan was found to exist in two main conformations: the two-fold (^2f^Xn) and three-fold (^3f^Xn) screw conformations. The ^2f^Xn conformation adopts a flat-ribbon shape, with a 360° rotation between every two xylose units, due to folding onto the surface of cellulose microfibrils^14,16–18^. In contrast, the ^3f^Xn conformation rotates 360° every three-xylose unit, forming a helical shape with increased mobility, important for forming the matrix and interacting with lignin^19–22^. Lignin is mainly composed of *p*-hydroxyphenyl (H), guaiacyl (G), and syringyl (S) units linked by ether and carbon-carbon bonds. It plays a crucial role in maintaining the hydrophobicity of lignified tissues, and serves as a barrier to enzyme accessibility, thereby hindering the utilization of lignocellulosic biomass for energy applications.

Despite significant advances, a key unresolved issue is that xylan binding to the surface of elementary cellulose fibrils, as discussed above, may increase fibril size, potentially leading to overestimates of chain numbers^23–25^. Fibril diameters have been observed between 2 to 5 nm, with high-resolution microscopy and electron tomography suggesting approximately 3 nm for each elementary microfibril^10,26–28^. The influence of mobile three-fold xylan on fibrils has been largely mitigated by ssNMR studies, which have shown that three-fold xylan does not significantly contribute to fibrillar structures^14,21,22,29^. However, the binding of well-structured two-fold xylan chains to cellulose microfibrils may induce changes in the conformations of surface and interior chains, leaving uncertainties regarding the fibril chain number, precise structure, and arrangement.

In this work, intact ^13^C-labeled wheat straw (*Triticum aestivum*) are analyzed using ssNMR^30,31^, small-angle X-ray scattering (SAXS), and wide-angle X-ray scattering (WAXS) techniques to determine the dimensions and arrangement of xylan-bound cellulose fibril. The combined input of various ssNMR techniques on the detailed composition and interactions of structural polymorphisms of cellulose, xylan, and lignin, as well as cellulose surface-to-interior ratios and hydration and dynamics profiles in the heterogeneous cell wall, lead to the hypothesis that rigid two-fold xylan, when bound to cellulose, functions as adjunct surface chains, contributing to the size and shape of the core-shell fibril observed by SAXS. We identify xylan-bound cellulose microfibrils composed of 18 glucan chains and up to four two-fold xylan chains closely structured along the fibril axis, which serve as a secondary docking site for lignin, following its predominant interactions with three-fold xylan. These findings provide new insights into the grass-specific plant cell wall structure at the molecular level and are anticipated to aid in the development of more efficient biomass deconstruction and conversion into high-value products.

## Results

### Polymorphic molecular structures in the cell wall of wheat straw

Uniformly ^13^C-labeled wheat straw, free from chemical treatments, was subjected to atomic-level analysis using ssNMR; therefore, the chemical structure and physical packing of the predominantly cell walls were preserved intact in the near-native state. A series of 1D ^13^C ssNMR spectra were utilized to rapidly assess the distribution of biomolecules across distinct dynamical regimes (**Figure 1a**). The ^13^C ssNMR spectra of polysaccharides (60 to 110 ppm) exhibited a large number of peaks originating from cellulose and xylan, with diverse conformational structures (**Figure 1b**). Cellulose signals, including β-1,4-glucan chains residing on the surface (s) or internal (i) domains of the microfibril, showed strong signals in the 1D ^13^C cross polarization (CP) spectrum that predominantly detect rigid molecules. It has been suggested that the interior cellulose predominantly adopts the *trans-gauche* (*tg*) hydroxymethyl conformation, differs from the surface cellulose, which contains predominantly the *gauche-trans* (*gt*) conformation, and to a lesser extent, the *gauche-gauche* (*gg*) conformation^32–34^. These various hydroxymethyl conformations are crucial to maintaining the hydrogen bonding network’s stability and determining cellulose’s chemical reactivity. Meanwhile, signals from xylan chains, including those in two- and three-fold conformations (Xn^2f^ and Xn^3f^) were relatively weak in CP but became stronger in the 1D ^13^C direct polarization (DP) spectrum with a short recycle delay of 2 s for selective detection of mobile molecules. Aromatic carbons from lignin were observed in the region of 120-160 ppm, showing consistent spectral patterns in two ^13^C DP spectra with short and long recycle delays, as well as their difference spectrum. This evidence indicates that the lignin components present in the rigid and mobile fractions share a similar composition and have nearly equal populations.

**Figure 1.**
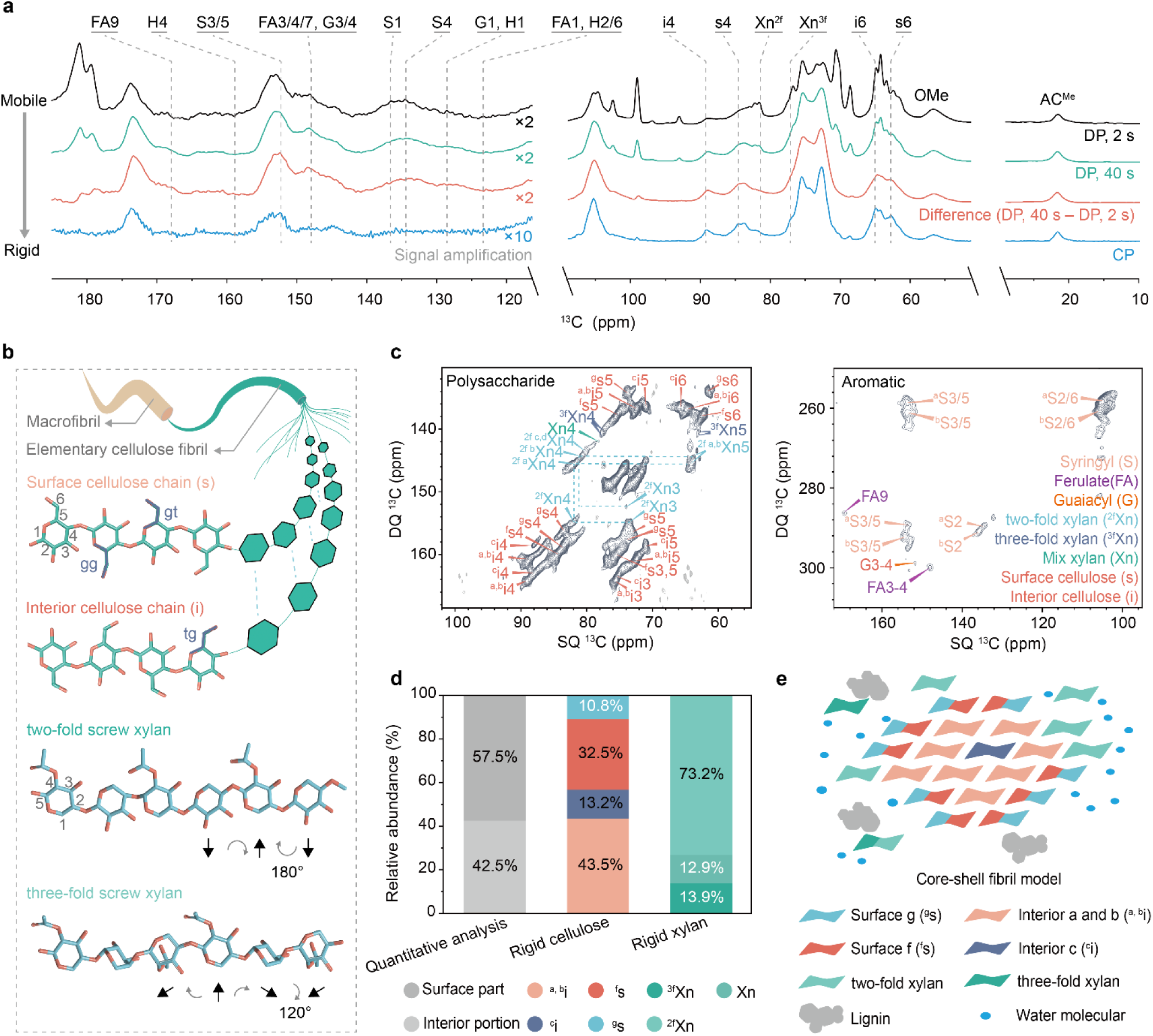
ssNMR reveals the polymorphic structure of molecules in intact wheat straw. **a,** Dynamic heterogeneity of polymers is revealed by 1D ^13^C ssNMR spectra. ^13^C Direct polarization (DP) enables selective detection of mobile molecules by employing short cycle delays, while Cross-Polarization (CP) selectively detects rigid molecules via ^1^H-^13^C dipolar couplings. Mobile molecules are preferentially tagged by DP 2 s and rigid components can be selectively observed using a difference spectrum and CP. **b,** Schematic description of representative structures of polysaccharides. The elementary cellulose fibrils is composed of surface cellulose chains and interior cellulose chains, which show distinct hydroxymethyl conformation (tg, gg, and gt). The two-fold conformation allows xylan to form a flat ribbon shape, rotating 360° every two xylose units. Xylan in the three-fold conformation rotates 360° every three xylose units, forming a helical shape. **c,** Representative 2D ^13^C CP J-INADEQUATE spectrum of wheat straw, elucidating carbon connectivity in rigid molecules. In this experiment, directly bonded nuclei are detected at the same double-quantum (DQ) chemical shift, which corresponds to the sum of their individual single-quantum (SQ) chemical shifts. Polysaccharide (left panel) and aromatic (right panel) signals are resolved using 2D ^13^C CP INADEQUATE methods. Abbreviations are given and molecules are color-coded (e.g., ^g^s5 denotes carbon 5 of glucose type-g; Xn4 denotes carbon 4 of two/three-fold mixed xylose). **d,** Contents of surface and interior cellulose domains. The quantitative analysis (grey; the first column) was achieved by deconvoluting quantitative 1D ^13^C DP spectra measured with long recycle delays of 40 s. The other two columns were determined using peak volumes of the rigid polysaccharides in 2D CP J-INADEQUATE spectra (**Table S2**). **e,** A possible cross-section model of xylan-bound cellulose microfibrils. The hydrophobic surface chains are labeled as ^g^s, hydrophilic surface chains as ^f^s, middle layer internal chains as ^a,b^i, and deeply embedded core chains as ^c^i.

The signals of rigid polysaccharides were assigned through a series of 2D ^13^C-^13^C correlation spectra (**Figure 1c** and **Table S1**). The structure of cellulose in wheat accommodates five types of glucose units, distributed as follows: residues embedded in the core (^c^i), those underneath the surface (^a^i and ^b^i), and those preferentially residing on the hydrophilic (^f^s) or hydrophobic (^g^s) surfaces^34,35^ (**Figure 1c**). The intricate nature of the hemicellulose structure is reflected in the observed peak multiplicity. Wheat xylan exhibited two-fold (^2f^Xn), three-fold (^3f^Xn), and mixed (Xn) conformations (**Figure 1c**). In the rigid fraction, around three-quarters (73%) are ^2f^Xn (**Figure 1d** and **Table S2**). Since the 2-fold flat-ribbon conformers are induced only when xylan folds onto the even surface of cellulose microfibrils, the uniquely high observed content is likely due to extensive xylan-cellulose interactions in wheat straw^14^. The 3-O-arabinose and glucuronic acid-substitutions of xylan chains were only observed in the mobile phase due to the flexibility of these sidechains (**Figure S1** and **Table S3**).

2D ssNMR and solution-state HSQC spectra revealed signals from syringyl (S), guaiacyl (G), and *p*-hydroxyphenyl (H) units within the aromatic domain (**Figure 1c** and **Figure S2**). H units are minimal at 0.5%, while S and G units dominate at 99.4%, emphasizing their structural prevalence. These monolignol units accommodate different numbers of methoxyl substitutions at the carbons 3 and 5 positions on their aromatic rings, leading to well-resolved signals in solid-state NMR spectra. The S units also exhibited two sets of signals due to their structural and conformational variations in solids.

High-resolution solid-state NMR data imposes three constraints on the average cross-sectional structure of xylan-bound cellulose microfibrils. First, analysis of 1D quantitative ^13^C DP spectra, measured with an extended recycle delay of 40 s, allows for peak deconvolution to determine the amount of the surface and interior cellulose domains, yielding values of 57.5% and 42.5%, respectively (**Figure S3** and **Table S4**). This value subsequently allowed the N_core_/N_elementary cellulose fibrils_ ratio to be extrapolated for counting the number of chains^29,36^. Second, analysis of peak volumes corresponding to five distinct cellulose conformers resolved in 2D spectra enables determination of the molar composition within rigid cellulose (**Figure 1d**). Third, cellulose microfibrils are ensheathed by xylan chains that adopt 2- fold, 3- fold, and mixed conformations in an approximate ratio of 5:1:1 in the rigid region. Integration of these findings reveals structural insights into the average cross-sectional architecture of cellulose, best fitting an 18-chain elementary microfibril with three flat-ribbon two-fold xylan chains near the microfibril surface and a three-fold xylan chain within a nanometer (**Figure 1e**).

### Dynamics and hydration profiles of biomolecules in the heterogeneous cell walls

To rationalize the location of fibrils and associated molecules within the native, heterogenous cell wall, we measured the dynamics and hydration of biomolecules using four ssNMR techniques. The dipolar order parameter (S_CH_) was determined using a dipolar-chemical-shift correlation (DIPSHIFT) experiment. This experimental scheme was carried out using two polarization approaches: CP for rigid molecules and 40 s DP for quantitative analysis of all molecules (**Figure S4**). Cellulose is intrinsically rigid, displaying large, near-unity S_CH_ values for i4 and s4 sites regardless of polarization methods, while xylan exhibits a broad range of dynamics, with large S_CH_ values of 0.90–0.95 in the rigid domain associated with cellulose and 0.7–0.8 for all xylans within the cell wall (**Figure 2a** and **Table S5**). This provides evidence of the dual function of xylan in both packing with cellulose microfibrils, thereby becoming rigidified, and in forming the mobile matrix.

**Figure 2.**
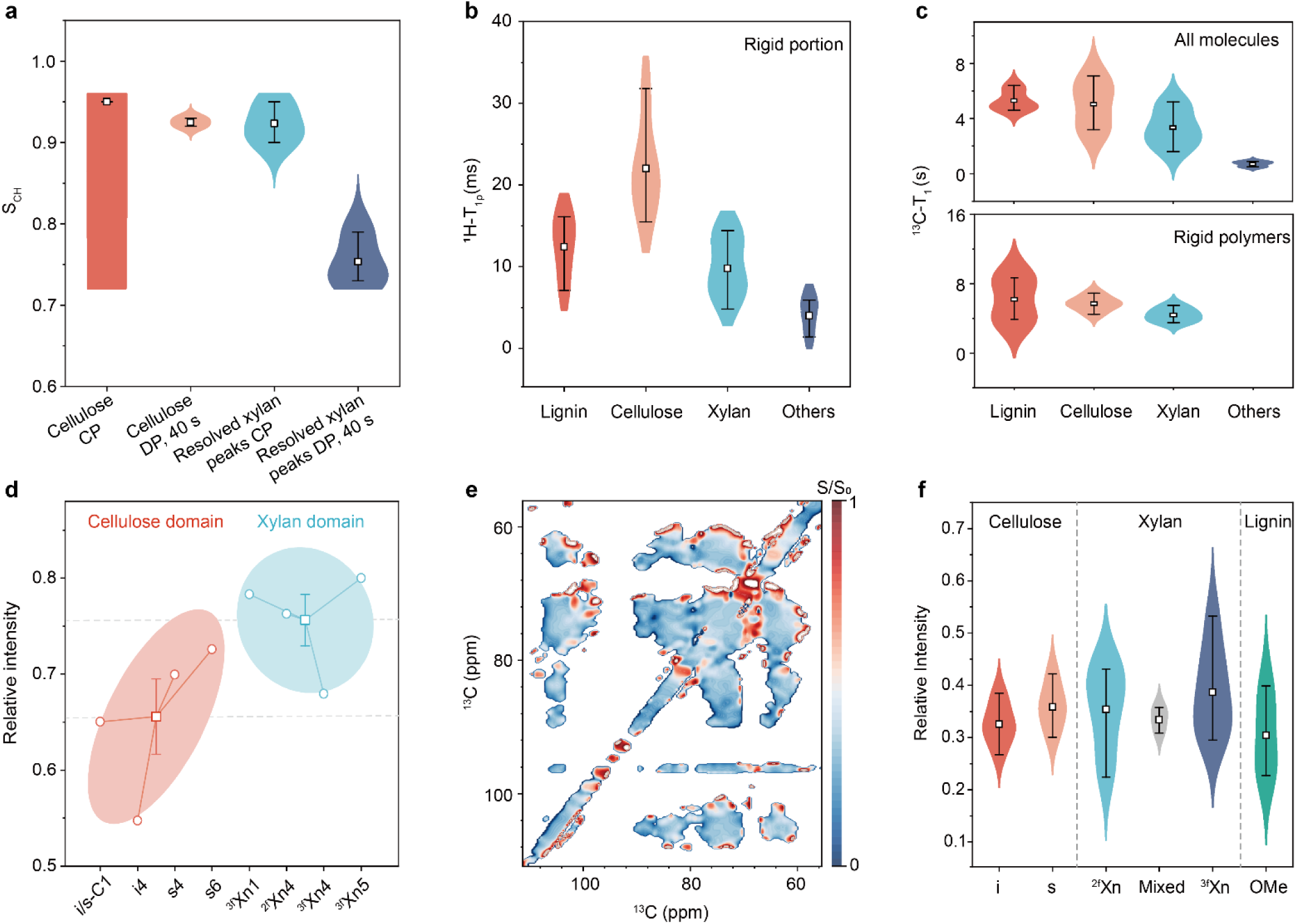
Carbon-specific hydration and dynamic landscape of polysaccharides and lignin. **a,** ^13^C-^1^H dipolar order parameters (S_CH_, 0 < S_CH_ < 1) of cell wall polysaccharides. n = 2, 2, 3, and 3 for CP cellulose (i4 + s4), DP 40 s cellulose (i4 + s4), CP resolved xylan peaks and DP 40 s resolved xylan peaks, respectively. The open box represents the mean and the error bars are s.d. **b,** ^1^H-T_1ρ_ relaxation times of cell wall polymers (n = 4, 4, 4, and 3 for lignin, cellulose, xylan, and other matrix carbohydrates, respectively; mean and s.d.). **c,** ^13^C-T_1_ relaxation times of all molecules (top panel; n = 6, 5, 5, and 5) through quantitative DP detection, and rigid molecules (bottom panel; n = 3, 5, and 5 for lignin, cellulose, and xylan, respectively) as indicators of nanosecond-timescale motions. **d,** Water ^1^H spin diffusion in cellulose and xylan, demonstrating the interaction of water with polysaccharides (mean and s.d.). **e,** Hydration intensity ratio (S/S_0_) plotted as a 2D spectral heat map in the polysaccharide region. **f,** The distribution of relative water-edited intensities (S/S_0_) of different carbohydrate and lignin forms and sites: n = 6, 5, 7, 3, 6, and 4 for i, s, ^2f^Xn, mixed, ^3f^Xn and lignin-OMe, respectively. The open box represents the mean and the error bars are s.d.

Consistently, xylan and other carbohydrates in the matrix had short ^1^H-T_1ρ_ relaxation time constants of < 15 ms (**Figure 2b** and **Figure S5**), indicative of the coordinated movement of their sugar rings. In contrast, the cellulose domain exhibits the longest ^1^H-T_1ρ_ relaxation times (15–30 ms) due to the constrained motions within the extensively hydrogen-bonded microfibrils. The ^1^H-T_1ρ_ time constants of the surface sites are 5-10 ms shorter than those of the interior sites (**Table S6**), indicating that the surface chains of cellulose microfibrils are more flexible than the interior chains^30,33,37^. The ^1^H-T_1ρ_ time constants of lignin fall between those cellulose and matrix polysaccharides, revealing its partial flexibility (**Figure 2b**).

With quantitative DP detection of all molecules, the ^13^C-T_1_ relaxation time constants decreased in the order of lignin, cellulose, xylan, and other matrix carbohydrate components (**Figure 2c** and **Table S7**). The relatively short ^13^C-T_1_ relaxation time constants of xylan and other matrix components emphasized their capabilities of accommodating rapid local motions on the characteristic nanosecond timescale. The long ^13^C-T_1_ relaxation time constants of lignin and cellulose carbon sites indicate that aromatics and cellulose microfibrils resist rapid local reorientation motions, consistent with previous findings in other grass species, such as *Zea mays*^21^. A similar trend was observed for the rigid molecules detected using CP, despite the loss of signals from the highly mobile components in the matrix. Notably, the ^13^C-T_1_ relaxation of lignin in the rigid fraction was highly heterogeneous, indicating that spin-exchange occurred between a portion of lignin and some polysaccharides in close spatial proxiimity^19,22^, while other lignin polymers remain within hydrophobic nanodomains^21,27^.

The hydration profiles of biomolecules were examined through water-editing experiments^38,39^, which showed preferential hydration of xylan over cellulose (**Figure 2d** and **Figure S6**). The hydration profiles were quantified by measuring the water-edited intensities (S/S_0_) of 31 carbon sites from various carbohydrates and lignin units, which were resolvable in a 2D hydration heatmap (**Figure 2e**, **Figure S7,** and **Table S8**). Among all polysaccharides, interior cellulose and mixed conformers of xylan exhibited the lowest average hydration levels (**Figure 2f**). The former is expected, as the core of each microfibril consists of relatively inaccessible cellulose, while the latter is unexpected and may suggest that the intermediate conformations between the two maxima of two-fold and three-fold xylan are caused by trapped xylan in partially dehydrated domains. This explains why the mixed intermediate conformations of xylan were most pronounced in fully dehydrated Arabidopsis and some wood stems^22,40^, where certain structural domains may be partially dehydrated, but were not observed in well-hydrated maize, rice, and switchgrass stems^21^. A slightly higher hydration level was observed for the surface chains of cellulose and the two-fold xylan chains, two molecules tightly packed together, exhibiting similar hydration levels. The highest hydration level was observed in three-fold xylan, consistent with the important role of this molecule in regulating water activity in the matrix. Meanwhile, lignin-OMe exhibits the lowest hydration level due to the hydrophobic nature of these aromatic polymers and their self-aggregation into nanodomains in the cell corners^21,41^.

### The patterns of biomolecules packing in the cell wall

To investigate polymer packing in lignified wheat cell walls, we measured 2D dipolar-gated proton-driven spin diffusion (PDSD) spectra with short (0.1 s) and long (1.0 s) mixing periods, the latter of which yielded 97 intermolecular cross-peaks on the sub-nanometer scale (**Figure 3a**). For example, magnetization transfers from the acetyl methyl of xylan (Ac^Me^; source) to site 3 or 5 of lignin (S3/5; sink) resulted in a cross peak at (21, 153) ppm. These intermolecular interactions are classified into five types based on the structural motifs of the contacting polymers: (i) cellulose and xylan acetyl groups, (ii) lignin and cellulose, (iii) different lignin units, (iv) lignin and xylan, and (v) lignin and mixed sugars (**Figure 3b, c** and **Table S9**). These intermolecular cross-peaks are further classified as strong, medium, or weak interactions based on their relative intensities.

**Figure 3.**
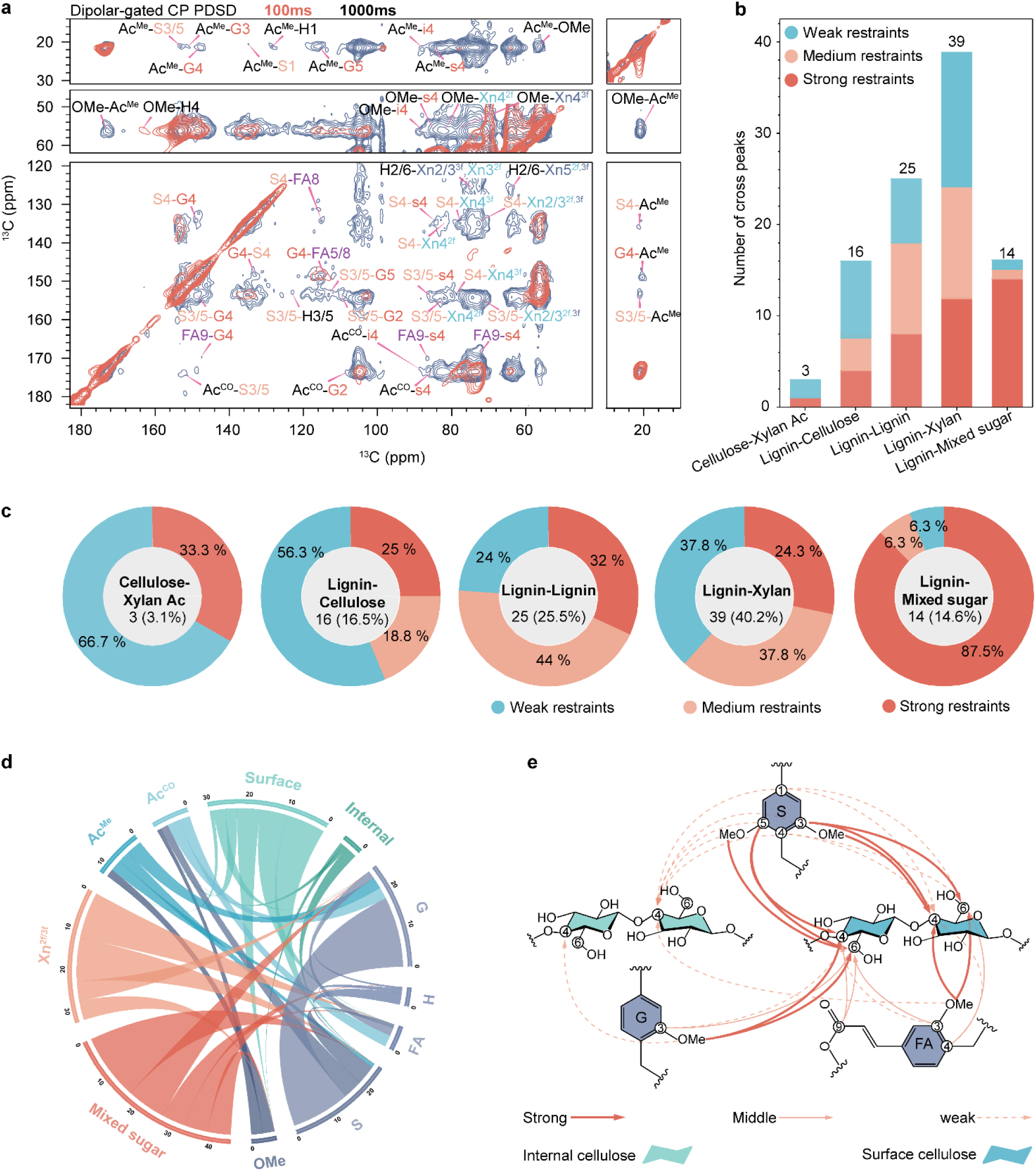
Detection of intermolecular interaction pinpointing packing interactions between polymers. **a**, Intermolecular cross peaks identified in dipolar-gated 2D ^13^C correlation spectra were detected with long (1000 ms PDSD) and short (100 ms PDSD) mixing periods. Abbreviations are given and molecules are color-coded (e.g., Ac^Me^-S3/5 denotes the source Ac^Me^ to sink S3/5.). **b**, Summary of polymer interactions. In 1D slices extracted from 2D gated PDSD spectra, peaks with intensities greater than 4% are classified as strong restraints (dark orange), those between 2% and 4% are considered medium restraints (light orange), and peaks with intensities below 2% are categorized as weak restraints (light blue). **c**, The percentage of strong, medium, and weak cross peaks within each interaction type (e.g., strong restraint accounts for 33% in of cellulose-xylan acetyl interactions). **d**, The interactions of molecules in the wheat straw were plotted. Each arc represents a source or sink, while the lines connecting the circles illustrate the interactions between the two components. The thickness of the line represents the strength of cross peaks. **e**, Structural summary of lignin-cellulose packing interactions. Arrows show the direction of polarization transfer from source to sink; Thick lines represent strong intensity, thin lines medium intensity, and dashed lines weak intensity.

The most extensive interactions were observed between lignin and xylan, with 39 cross peaks (**Figure 3b**; **Figure S8** and **Table S9**), 62% of which are either strong or medium cross peaks (**Figure 3c**). This supports xylan’s predominant role in initially anchoring and subsequently packing with lignin in the secondary cell wall. More interestingly, lignin, particularly its OMe units, exhibited greater spatial proximity to two-fold than three-fold xylan (**Figure S8b**), suggesting that distinct xylan conformers mediate different functional roles in lignin. In line with recent computational studies, some two-fold flat-ribbon xylan chains orient their acetyl groups toward lignin when adsorbed onto the cellulose surface^19^. The next most dominant interactions occur between different monolignol units, with 25 cross-peaks observed, 76% of which are strong. This is expected, as these units are covalently linked to form a three-dimensional heterogeneous, amorphous and hydrophobic polymer network.

Lignin also exhibited correlations with cellulose as a secondary interaction site, with 16 cross-peaks observed (**Figure 3b**). However, most (56%) of these cross-peaks were very weak, indicating that the average spatial proximity of cellulose-lignin interactions is lower than that of cellulose-xylan (**Figure 3c**). This observation aligns with recent findings that cellulose-lignin interactions typically occur in crowded systems, particularly in mechanically strong woody stems^22^.

A statistical summary of the quantity and strength of intermolecular interactions highlights the critical role of carbonyl and methyl carbons in acetyl groups (Ac^CO^ and Ac^Me^) in stabilizing xylan’s interactions with other molecules (**Figure 3d**). These key carbon sites exhibited strong interactions with methoxy (OMe) and ring carbons of S and G units of lignin, as well as the surface chains of cellulose microfibrils. S units also showed multiple correlations with surface cellulose, while G units did not, indicating that the additional methoxy group increased the tendency of establishing physical contact between the lignin and the fibrillar surface^42^.

### The fitting geometry and chain number calculation of xylan-bound cellulose fibrils

SsNMR data have been employed to estimate the fibril size through the integration of data on structural polymorphism and surface-to-interior ratio using simplified core-shell elementary cellulose fibrils models^17,21,22,29^. X-ray scattering techniques, such as SAXS and WAXS, were also employed to characterize the average nanoscale geometry of the fibril. We combined these complementary methods here to constrain the size of fibrils and special conformation of glucan chains, proposing a structural view that additional two-fold xylan chains bind to elementary cellulose fibrils to form xylan-bound cellulose microfibrils. Subsequently, the potential cellulose arrangement habits and chain number of the xylan-bound cellulose microfibrils were further proposed.

The 2D SAXS pattern and 1D fitting curves display a standard pattern of lignocellulosic biomass with anisotropic characterization (**Figure 4a, b**). Although the features of fibrils in the SAXS pattern are dominated by the form factor (size and shape of fibrils) and structure factor (the arrangement of fibrils), the surrounding mobile matrix likely influences the estimation of fibril size due to strong interactions. To precisely measure the size of the fibril cross-section, SAXS data were used to model the entire fibril using porod analysis, which suggested a relatively sharp interface between fibrils and the matrix (**Figure S9a–d**). The fibril model, represented by circular cylinders (CYL), was fitted to determine the fibril geometry using SASview software with WoodSAS (**Figure S9e** and **Table S10**). The cross-sectional radii of gyration (Rc) and fibril radius in the isotropic sample exceeded those in the anisotropic sample, suggesting that fibril-matrix interactions may contribute to geometry and an overestimation of cross-sectional areas (**Figure 4c–d**). Consistent with ssNMR observations, SAXS provides additional insights for guiding the construction of a representative packing structure of the cell wall. Under fully hydrated conditions, the SAXS intensities reflect the stronger characteristics of the structure factor due to the increasing distance (a = ∼27 Å) among fibrils, which can be easily changed because of a lower packing of the cellulose microfibrils. These results suggest that cellulose microfibrils in the cell wall are separated by readily hydrated polysaccharides, which is consistent with our ssNMR data and previous studies^25,43^. When the sample is hydrated, the water molecules can easily infiltrate into the spaces between the cellulose microfibrils, leading to significant swelling^44^. Additionally, the measured average lattice plane spacing of the crystalline were 0.61, 0.53, and 0.43 nm which corresponds closely to cellulose Iβ—0.61/0.54/0.40 nm along the directions of (1– 10)/(110)/(200), respectively (**Figure 4e–g**; **Table S11**). The cellulose Iβ part may contribute from core cellulose, in which its presence can be detected by ssNMR and X-ray scattering (**Figure S10a–c**).

**Figure 4.**
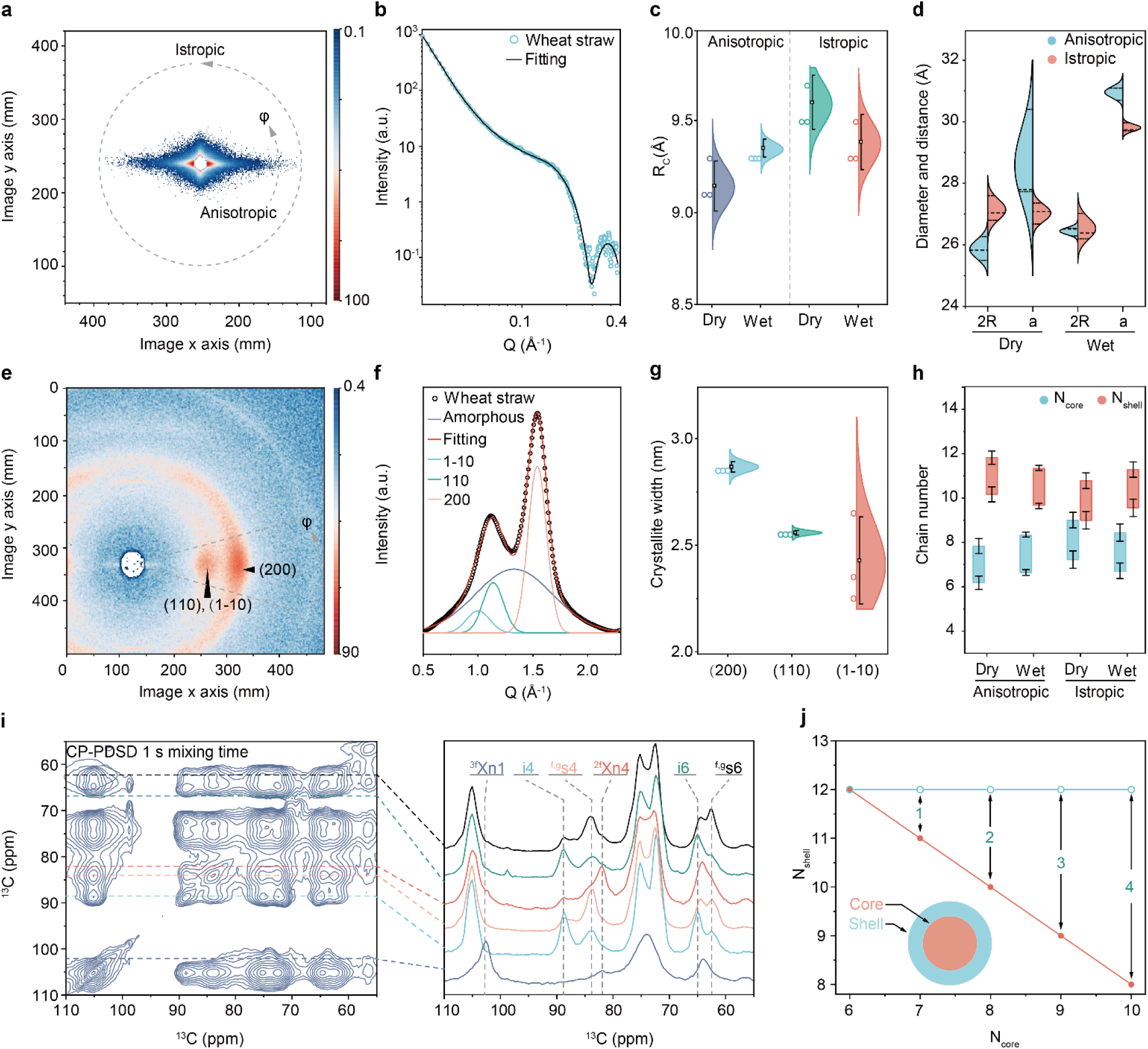
Potential architecture of xylan-bound cellulose microfibrils. **a,** Two-dimensional SAXS pattern of the wheat straw. When subjected to an X-ray beam, the pattern reveals a distinctly anisotropic scattering, indicative of nanoscale fibril structures. The anisotropic (∼25°) and isotropic (full azimuthal 360°) 1D profiles were extracted from 2D SAXS. **b,** Anisotropic integration of the SAXS profile was fitting using an assembly of hexagonally packed cylinders model ^50^. **c,** The cross-sectional radii of gyration (R_c_) and **d,** Diameter (2R) of fibril and distance (a) among fibrils were plotted (Relative fitting parameters: n = 3 for all samples; mean and s.d.). **e,** Two-dimensional WAXS patterns of the wheat straw. **f,** The peak profile is integrated by software fit-2D (Equatorial direction) and analyzed by deconvolution. **g,** Crystallite widths, measured from fitting parameter (Crystalline width: n = 3 for 200, 110, and 1-10, respectively; mean and s.d.). **h,** The estimated cellulose chain number of core-shell fibril was calculated using SAXS fitting parameters and ssNMR data (Chain number: n = 3 for all samples; mean and s.e.). **i,** Left-hand side: CP-PDSD 1 s mixing time ssNMR spectrum of the wheat straw. Right-hand side: 1D slices extracted from the 2D pattern. Each 1D slice and its corresponding diagonal peak are in the same color, e.g. black for ^f,g^S6 in the top slice and light cyan for ^2f^Xn5 and the second slice. **j,** Based on the assumption of 18 cellulose chains, the numbers of surface and core chains in xylan-bound cellulose microfibrils were defined for interpretation. e.g., The result (N_core_ = 8; N_shell_ = 10) indicates that 8 cellulose chains are forming a crystalline core, while 12 surface chains, along with 2 additional xylose chains, constitute the surface layer that surrounds and encapsulates this core.

The core cross-section area and chain number of fibril can be deduced assuming a shell thickness was 4–5 Å, supported by previous studies^32,37^. The calculated mean chain number of the interior crystalline core (N_core_) ranges from 6 to 8 chains (2.0–2.7 nm^2^). Employing N_core_ values derived from SAXS parameters, the estimated R values for 18-, 24-, and 36-chain models were 34–43%, 26–33%, and 17–22%, respectively. The experimentally measured R (∼42.8%) closely aligns with the 18-chain model’s predicted range (34–43%), reinforcing the validity of the 18-chain assumption. Based on the 18-chain model assumption, the surface chain number was estimated at 9 to 11 chains (**Figure 4h**). Interestingly, the inferred mean core chain number of 8 (∼42.8% corresponds to 8 core chains) exceeds the typical 6-core chain count. We propose that two additional surface chains are required to accommodate the potential fibril habit (234432) thereby maintaining structural integrity (**Figure S10d**). To meet the 18 chains (with 8 core chains), there would be more additional surface chains to meet the fibril habit (8 core + 10 surface chains + additional surface chains).

To explore the extra chain number in fibril, we hypothesize that the additional chains consist of well-structured two-fold xylan, which is closely bound with elementary cellulose fibrils based on the 18-chain model. To assess the involvement of xylan in fibril structure, we measured 2D PDSD spectra with a long mixing time (1 s) to probe intermolecular interactions happening between different carbohydrates (**Figure 4i** and **Figure S10d**). The observed intermolecular cross peaks represent close spatial proximity between xylan and both the internal glucan chains of cellulose (e.g., ^2f^Xn4-^a, b, c^i6/4) and surface chains of cellulose (e.g., ^2f^Xn4-^f, g^s4/6). The results showed that xylan’s interactions with internal and surface cellulose chains were strong for ^2f^Xn, as shown by the strong cellulose peaks in the ^2f^Xn4 cross-section, but much weaker for ^3f^Xn, as shown by the weak cellulose peaks in the ^3f^Xn1 cross-section. Contrary to the widely accepted view that primarily glucan chains define the cross-section area of the fibril, our findings indicate that both glucan and two-fold xylan chains significantly contribute to the fibril’s section area. Consequently, these structures are termed xylan-bound cellulose microfibrils.

Integrating SAXS data and ssNMR results, we propose that there is substantial contact between both the surface and interior residues of the fibril and a considerable number of xylan chains. These xylan chains likely reside on cellulose surfaces, aligning with the observed fibril habits. Additionally, it will be another alternative to match the fibril model, representing an average result based on 8 core chains in the complex cell wall environment. Each xylan-bound cellulose microfibril comprises a minimum basic unit of 18 glucan chains within its cross-section. As the core chain number increases, the number of additional xylose chains may range from 0 to 4, thereby accommodating various glucose residues in different fibril habits (**Figure 4j** and **Figure 5a**). However, we cannot exclude that more than 4 two-fold xylan chains may be present in some segments of the fibril. Using SAXS and ssNMR data, we have fitted the chain number of xylan-bound cellulose microfibrils, yielding interpretations that support the presence of averaged 18 glucan chains, variably associated with 0–4 two-fold xylan chains closely aligned with the cellulose core in the different parts of fibrils. This interpretation accounts for the molecular interactions between xylan and cellulose and suggests that the conformation of surface glucans on xylan-bound cellulose microfibrils may adopt an interior-like conformation to reconcile the ratio (R) and geometry, corresponding to the previous research ^14,19,35,45–49^.

**Figure 5.**
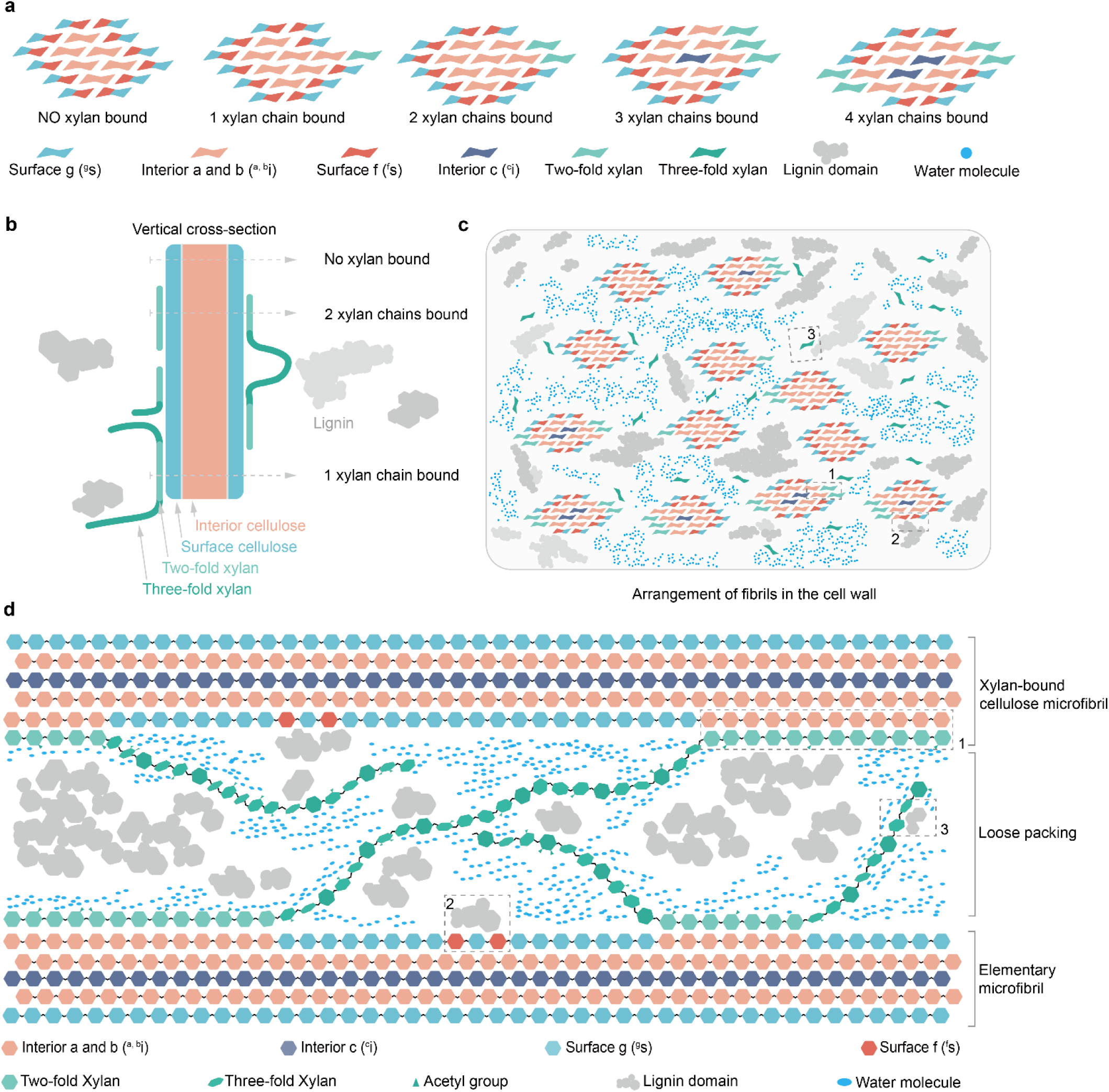
Conceptual diagram of xylan-bound cellulose microfibrils in the cell walls of wheat straws. **a,** The cross-sectional view of xylan-bound cellulose microfibrils shows the detailed possible conformations. For example, an 18-chain fibril has 6 interior chains and 12 surface chains when no xylan chains are bound or has 7 interior chains and 11 surface chains when 1 xylan chain is bound. **b,** The vertical-sectional view of a xylan-bound cellulose microfibril displays a different cross-sectional view. **c**, The xylan-bound cellulose microfibrils of different habits arrangement in the cross-sectional view. **d**, This diagram presents the spatial organization of lignin, cellulose, two-fold xylan, three-fold xylan, and water molecules in wheat straw’s cell wall. The numbered labels indicate specific structural features: (1) two-fold xylan binds with the cellulose surface, which alters the surface conformation to the interior conformation; (2) the Lignin domain was in contact with the cellulose surface; (3) the Lignin domain was contacted with three-fold xylan. (4) The loose packing among xylan-bound cellulose microfibrils. The illustration considers the molecular fractions of polysaccharides, though it may not be precise to scale.

## Discussion

This study presents an alternative molecular model of xylan-bound cellulose microfibril architecture in wheat plant cell walls through the integration of ssNMR and X-ray scattering techniques. When fitting the data to an 18-chain elementary cellulose fibril model, a variable number (0–4) of two-fold xylan chains appears closely aligned with the cellulose microfibrils (**Figure 5a**). Previous ssNMR and X-ray diffraction studies have sometimes favored a canonical 24-chain fibril model or binding arrangement^33,37^ (**Figure S11**). However, it is important to recognize that these structural interpretations represent averaged models that inherently account for cellulose coalescence, where multiple elementary microfibrils merge in specific regions, likely forming larger crystalline bundles that increase the apparent size of cellulose and contribute to the observed larger dimensions (**Figure S11**)^26,27,46^. Recent insights from high-resolution and cryogenic electron microscopy, scattering techniques, ssNMR, and modeling of the cellulose synthase complex, along with cellulose microfibrils synthesized *in vitro* and located *in muro* increasingly converge to support an 18-chain elementary microfibril model^6,9–12^. Our analysis suggests that even a simple 18-chain arrangement (234432) can sufficiently meet the necessary criteria whereas alternative models (34443 and 33333) are less compatible with the observed size and glucose residue environment, consistent with previous quantum modeling results^51^.

As the xylan-bound cellulose microfibril model is larger than the 18-chain model derived solely from SAXS results^46^, the excessed width in the cross section should be attributed to the incorporation of additional two-fold xylan chains^17,47^. These chains, aligning with the conformation of glucan chains on the cellulose microfibril surface, may contribute to an expanded size and occupy a significant portion of the elementary cellulose fibrils surface^52,53^, as they exhibit similar average dynamics and hydration (**Figure 2f**). A possibility is that the cross-section of a segment of the xylan-bound cellulose microfibril can commonly combine with an average of 2 chains of two-fold xylan (**Figure 5b**), but local variations may occur, ranging from 0 up to 4 along the fibril axis (**Figure 5a, b**). While this model is predominantly based on individual xylan-bound cellulose microfibrils, it does not preclude that cellulose coalescence may occur along portions of their length in wheat cell walls, although the formation of larger macrofibrils is more frequently observed in woody plants^54^. When coalescence occurs, the number of bound xylan chains is simultaneously reduced to achieve a good fit with our results. Notably, the presence of deeply embedded core chains, referred to as interior c (^i^c) in **Figure 5a**, can be accommodated when the number of cellulose-bound chains is 3 or 4, and this accommodation is further supported when coalescence occurs. Our updated model fits ssNMR and diffraction results align with recent in silico predictions and simulations^9,55–57^, and demonstrates how these views of averaged cellulose structure can meticulously be integrated with the characteristics of the 18-chain elementary microfibril synthesized by the cellulose synthase complex^6^.

To reconcile the two-fold xylan-bound cellulose model with the observed surface-to-interior domain ratios, some surface glucosyl residues should undergo hydroxymethyl conformational adjustments, shifting from *gt/gg* to *tg*, upon xylan binding, thereby transforming into “interior-like” domains (**Figure 5** and **Figure S12**). Our results also support the hypothesis that two-fold xylan binding to the cellulose surface increases the size and rigidity of xylan-bound cellulose microfibrils, thereby contributing to their recalcitrance^24,49,55^. Although there is ongoing debate regarding the conformational distribution of surface and interior cellulose chains, most experimental data suggest that surface chains primarily adopt the *gg*/*gt* conformation, with recent ssNMR measurements supporting the dominance of the *gt* conformation on the surface, while the *tg* conformation, stabilized by hydrogen bonding, is typically found in interior chains^34,58^. At this moment, it remains an open question how the structural rearrangement of hydroxymethyl conformation due to xylan binding, and potentially further influenced by cellulose coalescence, is facilitated in *muro*.

Our ssNMR data showed that lignin was primarily associated with three-fold xylan, while two-fold xylan and its bound surface cellulose acted as secondary interactors (**Figure 5c, d**). While the composition and structure of lignin polymers are plant-specific, the underlying physical principles regulating their interactions with carbohydrates are universal, and consistently observed in this study and many others across various plant species^16,17,22^. In wheat, lignin domains can be considered as two distinct parts due to their even distribution between rigid and mobile regions: one part of lignin may assemble into nanodomains to repel water, consistent with previous work^59^, while the remaining lignin interacts with polysaccharides in the secondary cell wall.

These findings, in conjunction with those of a recent study on biomass, indicate that the extensive interaction among biopolymers results in interentanglement^19,21,22^. In previous studies, hemicellulose is primarily responsible for coating cellulose microfibrils, followed by lignin^60^, and both simultaneously align parallel to the fibril axis^61^. Our ssNMR data on wheat straw further reveal the abundance of xylan-lignin and cellulose-lignin interactions (**Figure 5c, d**), where direct physical contact between the cellulose surface and lignin, along with the association between xylan and lignin through covalent linkages or physical packing, may reduce enzyme accessibility and contribute to biomass recalcitrance^62,63^.

SAXS and ssNMR results indicate that xylan-bound cellulose microfibrils are loosely packed in the cell wall, allowing water molecules to infiltrate the interfibrillar spaces. Meanwhile, interactions among elementary microfibrils, which are essential for maintaining cell wall mechanics^13^, and interactions between microfibrils, xylan, and lignin are stabilized through both covalent bonds and non-covalent interactions, contributing to the structural integrity of the plant cell wall. Water accessibility in the plant cell wall can influence the efficiency of enzymatic hydrolysis during biofuel production, as well-hydrated and loosely packed xylan-bound cellulose microfibrils are more susceptible to enzymatic attack. Conversely, the enhanced rigidity and reduced accessibility in less hydrated regions may contribute to biomass recalcitrance, presenting challenges for its conversion into biofuels and other value-added products^5,64^. Therefore, Understanding the architecture of xylan-bound cellulose microfibrils can aid in developing plant traits to overcome lignocellulosic biomass recalcitrance, improving biofuel and biomaterial production^65,66^. Future research should focus on identifying the molecular drivers of biomass deconstruction to improve biomass utilization^67^, further advancing our ability to effectively manipulate and use plant biomass.

## Methods

### Preparation of ^13^C-labelled wheat straw

Uniformly ^13^C-enriched wheat straw (U-60416, 95 atom% ^13^C) was obtained from IsoLife bv (Wageningen, The Netherlands), cultivated from seed to harvest under carefully controlled conditions. In brief, wheat plants (*Triticum aestivum*) were uniformly labeled with ^13^C to achieve an isotopic enrichment of 95 atom% at 16 weeks post-sowing. Plants were cultivated hydroponically in custom-designed, airtight labeling chambers capable of maintaining a closed atmosphere and high-irradiance conditions. The cultivation environment featured a photosynthetic photon flux density (PPFD) of 800 μmol m^-2^ s^-1^ at plant canopy height, with a photoperiod cycle of 16 hours light and 8 hours darkness. Ambient temperatures within the chambers were maintained at approximately 24 °C during the day and 16 °C at night, with relative humidity levels set at 75% during daylight and 80% overnight. A stable concentration of ^13^CO (average concentration of 400 ppm during the illumination period) was maintained by regulated injection from pressurized gas cylinders. Nutrients and water were supplied through aerated Hoagland-type solutions supplemented with essential micronutrients and iron, with nitrogen concentrations kept within the range of 25–200 mg L^-1^, pH values between 5.0 and 6.5, and electrical conductivity (EC) maintained between 0.4 and 0.7 mS cm^-1^. Immediately following harvest, plant tissues were carefully separated; leaves and stems were sectioned into smaller fragments and rapidly stored at -30 °C prior to freeze-drying (lyophilization). Post-lyophilization, samples were preserved under dark and dry conditions at 18 °C. The wheat straw was hydrated to approximately 40 wt%, after which 30 mg of the material was cut into slices and loaded into a 3.2 mm Bruker MAS rotor for ssNMR experiments ^68^.

### Solid-state NMR analysis of wheat straw

Solid-state ^13^C MAS NMR experiments on wheat straw samples were performed using a 3.2 mm Bruker E-free probe on a 800-MHz Bruker Avance Neo spectrometer. The experiments were carried out at a temperature of 295 K with a magic angle spinning (MAS) frequency of 19 kHz. ^13^C chemical shifts were externally calibrated using the CH_2_ signal of adamantane, set at 38.48 ppm on the tetramethylsilane (TMS) scale. Typical radio-frequency field strengths employed during the experiments ranged from 62.5 to 83.3 kHz for ^1^H and 50 to 71.4 kHz for ^13^C.

2D CP refocused J-INADEQUATE spectra were utilized for resonance assignment of polysaccharide and lignin signals. This technique selectively highlights cross-peaks between single quantum (SQ) and double quantum (DQ) chemical shifts of two covalently bonded carbons^69,70^. The DQ chemical shift represents the sum of the SQ shifts of two bonded carbon atoms, enabling enhanced resolution of overlapping signal pairs in two-dimensional space. Signal assignments were conducted concerning the Complex Carbohydrate Magnetic Resonance Database (CCMRD) and pertinent literature ^71^. Specialized solid-state NMR techniques were employed to selectively detect rigid and mobile molecules in cell wall materials, based on their inherent dynamic properties. The ratio of polysaccharides was determined by analyzing the volumes from well-resolved peaks in the 2D spectra. ^13^C direct polarization (DP) with a short recycle delay of 2 s was utilized to preferentially detect mobile molecules that have rapid ^13^C-T_1_ relaxation. To quantitatively probe all polymers, a longer recycle delay of 40 seconds was used for the ^13^C DP experimental scheme.

### Solid-state NMR analysis for structure

Intermolecular contacts were identified using 2D ^13^C-^13^C CP-PDSD with a mixing time of 1000 ms. To enhance the detection of lignin signals in the presence of proton-rich polysaccharides, ^13^C-^1^H dipolar coupling was reintroduced using a gated decoupling period in a modified proton-driven spin diffusion (gate-PDSD) experiment, with PDSD mixing times ranging from 100 ms to 1000 ms ^72–74^. This dipolar dephasing period was asymmetrically placed relative to the π pulse in the Hahn echo sequence to maximize the dephasing, incorporating two uncoupled delays of 32 μs and 16 μs. Cross peaks of wheat straw were identified and categorized into strong, medium, and weak restraints based on the relative intensities of each single cross peak within one-dimensional slices. Intensity cutoffs were established as follows: strong restraints (> 4.0%), medium restraints (2.0–4.0%), and weak restraints (< 2.0%). To visualize the molecular interactions, we constructed a chord graph. Restraints were classified into three categories— strong, medium, and weak—and assigned values of 3, 2, and 1, respectively, to represent their intensity in the graph.

We assessed the site-specific hydration profiles of polysaccharides using water-edited experiments, which provided measurements of hydration intensities^38,39^. The experiment employed a ^1^H-T_2_ relaxation filter to suppress the polysaccharide signals to < 5% of the original intensity, while retaining the majority of water magnetization (> 85%). The polarization of ^1^H from water was further transferred to nearby molecules using a 4 ms ^1^H mixing period, followed by a 1 ms cross-polarization (CP) step for ^13^C detection. Both the water-edited spectrum and the control spectrum were acquired with a 100 ms DARR mixing period. Water-to-polysaccharide/lignin one-dimensional buildup curves were generated using a ^1^H-T_2_ filter of 120 μs × 2, along with ^1^H mixing times ranging from 0 to 100 ms^38^. These S/S_0_ intensity ratios, which compare the intensities between the water-edited spectrum (S) and the control spectrum (S_0_), indicate the degree of water retention around various carbon sites in the cell wall.

To comprehensively characterize the molecular dynamics, we generated 63 relaxation curves by measuring ^13^C spin-lattice (T_1_) relaxation and ^1^H rotating-frame spin-lattice relaxation (T_1ρ_) relaxation. The ^13^C-T_1_ relaxation was measured using two methods: the CP-based Torchia T_1_ technique and the standard DP-based inversion recovery method^75^. Utilizing a z-filter, the Torchia-CP method preferentially detected rigid molecules in its z-filtered versions, while the inversion recovery experiment with a recycle delay of 30 s was used to quantitatively probe all molecules. Relaxation time constants were determined by fitting using single exponential functions. We measured the dipolar order parameter (S_CH_) using the dipolar-chemical-shift (DIPSHIFT) experiment at 7.5 kHz MAS. ^1^H-T_1ρ_ relaxation was detected using 62.5 kHz for the Lee-Goldburg spin-lock sequence^76^, with the scaling factor confirmed to be 0.577. CP-DIPSHIFT experiments were performed with a recycle delay of 2 s, while quantitative DP-DIPSHIFT experiments were conducted with a recycle delay of 20 s.

### Solution-state NMR experiments

Approximately 50 mg of ^13^C-labeled wheat straw was cut into small pieces and then milled into powder. Subsequently, approximately 20 mg of the sample was directly dissolved in 0.5 ml of dimethyl sulfoxide-*d_6_*(DMSO-*d_6_*). The dissolved sample was directly transferred into a 5 mm NMR tube for solution-NMR analysis. The 2D ^1^H-^13^C HSQC spectra were conducted at 289.2 K on a Bruker AVIII 400-MHz spectrometer equipped with a 5 mm BBO probe. The HSQC pulse program was based on published research, with parametric modifications.

### X**-** ray diffraction and WAXD data analysis

XRD data were collected at Xeuss 2.0 equipped with a Pilatus 3R 300K detector. A thin piece of wheat straw sample (5 mm × 5 mm × 0.538 mm) was affixed and attached to the sample holder. After a 300-second exposure, the Pilatus 3R 300K detector captured the signal and converted into two-dimensional patterned images. The images were converted into a 1D profile using fit2D software with a cake-type integration pattern. The 1D WAXS profile was fitted for 1–10, 110, and 200 peaks. The crystalline structure size (L) was estimated using the Scherrer equation: L = Kλ/(Bcos*θ*), where K is the form factor and is set as 0.90 according to the previous literature ^57,77^, λ represents the X-ray wavelength, B is the full width at half maximum (FWHM) of the diffraction peak measured in units of 2*θ*, and *θ* is the Bragg angle.

### Small-angle X-ray scattering and data analysis

SAXS data were collected at Xeuss 2.0 equipped with a Pilatus 3R 300K detector. The sample was fixed in a hole holder with an exposure time of 300 s. The resulting two-dimensional SAXS images were integrated to obtain the SAXS profiles (processing by Fit 2D), which were subsequently modeled using SasView software (version 5.0.6). SasView is a powerful tool to analyze SAXS, supporting a series of models.

The Porod plot (Q^4^ × I versus Q^4^) was analyzed. This approach was facilitated by the WoodSAS plugin available within SasView, as detailed in the software documentation^50^ (http://marketplace.sasview.org/). In a simplified representation, the distance (a) between the centers of adjacent fibrils and the cylinder radius (R) were determined. The cross-sectional radius of gyration (R_c_) is one of the most significant parameters in SAXS analysis ^37^. For example, the radius of gyration for a CYL with radius R is R_g_^2^ = R^2^/2. The chain number (N) can be calculated from the core cross-section area (S) ^33^: N = S/0.32. This calculation is based on the Iβ crystalline structure of cellulose, which is predominant in higher plants, where each cellulose chain occupies an approximate cross-sectional area of ∼0.32 nm² following previously established protocols^78^.

### Fourier Transform Infrared Spectroscopy

FTIR spectra were collected using a Nicolet iS10 FTIR-ATR spectrometer, with a resolution of 4 cm^−1^ across 16 scans, covering the spectral range from 4000 to 650 cm^−1^ range. To achieve full saturation with D_2_O, the wheat straw slice was presoaked in D_2_O for 24 hours.

## Supporting information

JACS Supporting information TW

## Acknowledgments

The authors express their gratitude for the financial support from the National Natural Science Foundation of China (Nos. 22178028 and U22A20422 to F. X.) and the Program of Introducing Talents of Discipline to Universities (Project 111, B21022 to F. X.). This research was supported by the ssNMR facility at the Institute of Drug Discovery Technology, Ningbo University, which assisted with data collection. A portion of the solid-state NMR analyses were supported by the U.S. Department of Energy under grant no. DE-SC0023702 to T.W. This work benefited from using the SasView application, originally developed under NSF award DMR-0520547.

## Author contributions

Conceptualization: Y.C.H., and F.X. Methodology: Y.C.H., S.X.Y., Z.L., P.X., Y.T.Z., G.H.M, Y.H., H.C.L., S.C., T.T.Y., T.W., and F.X. Investigation and formal analysis: Y.C.H., P.X., T.W., and F.X. Writing (original draft): Y.C.H., and F.X. Writing (review and editing): Y.C.H., P.X., T.W., and F.X. Funding acquisition: T.W. and F.X. Supervision: T.W. and F.X. Project administration: F.X.

