## Supplementary material for "Molecular Insights into Cell Wall Architecture and Xylan-Bound Cellulose Fibrils in the Wheat Straw": JACS Supporting information TW

Yucheng Hu *et al.*

*Corresponding authors:

 (T. Wang); (F. X.)

**This PDF file includes:**

Figure S1 to S12

Table S1 to S11

References (1 to 21)

### Figure

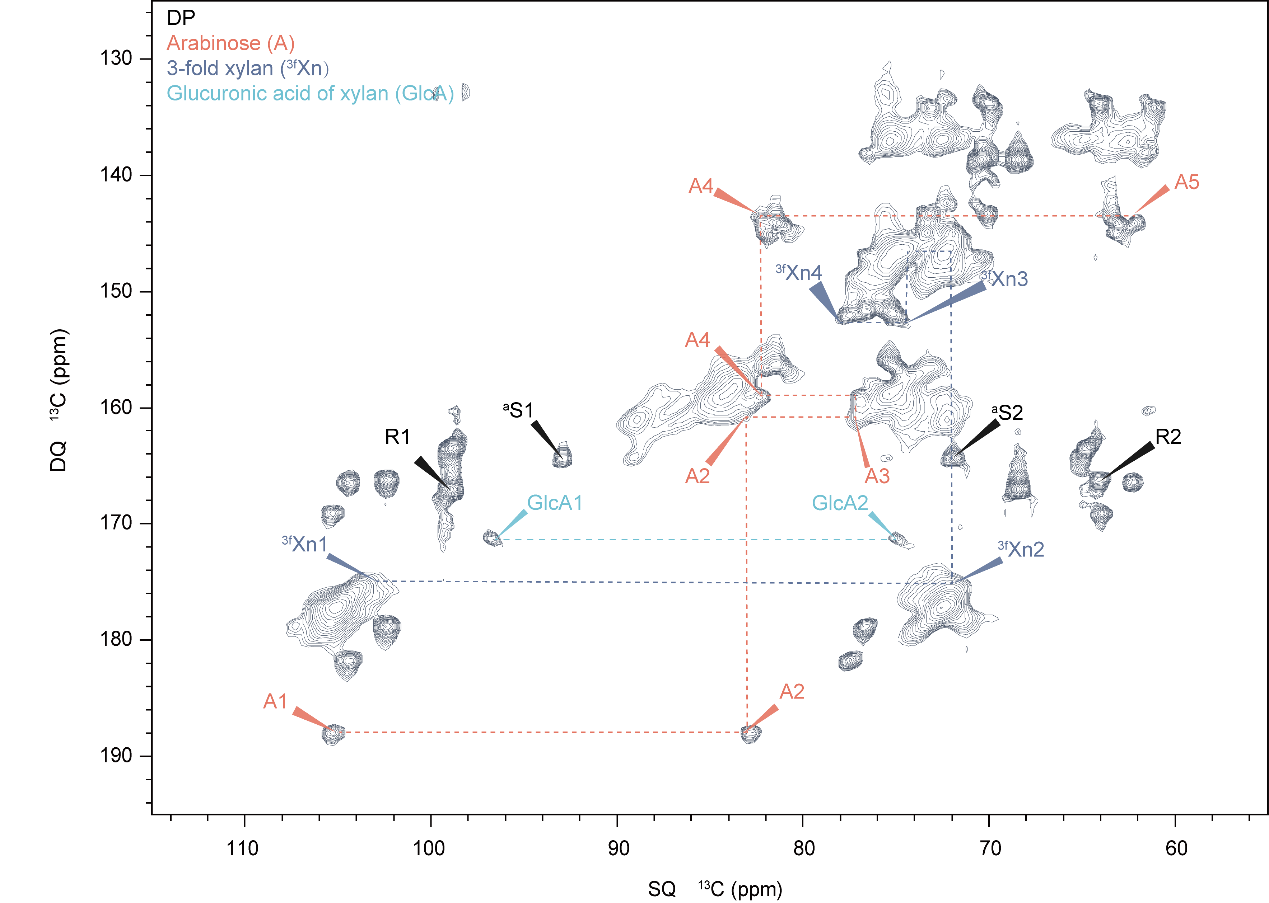

Figure S1. Mobile polysaccharides detected by refocused ^13^C DP-INADEQUATE experiments. Representative 2D ^13^C DP J-INADEQUATE spectrum of wheat straw, preferentially elucidating carbon connectivity in mobile molecules by employing short-cycle delays. By selectively highlighting the interactions between carbons in the mobile fraction, this technique is well-suited for distinguishing between the more rigid, structural components of lignocellulosic biomass (like cellulose) and the more flexible, easily exchangeable polysaccharides (such as hemicellulose). The mobile polysaccharides seem to be primarily found in the amorphous regions, where the polysaccharide chains are less constrained by crystalline cellulose or lignin networks. These regions contribute to important biochemical and physicochemical properties, such as enzymatic degradation and interaction with water, making them of significant interest in both fundamental research and applied biotechnological processes.

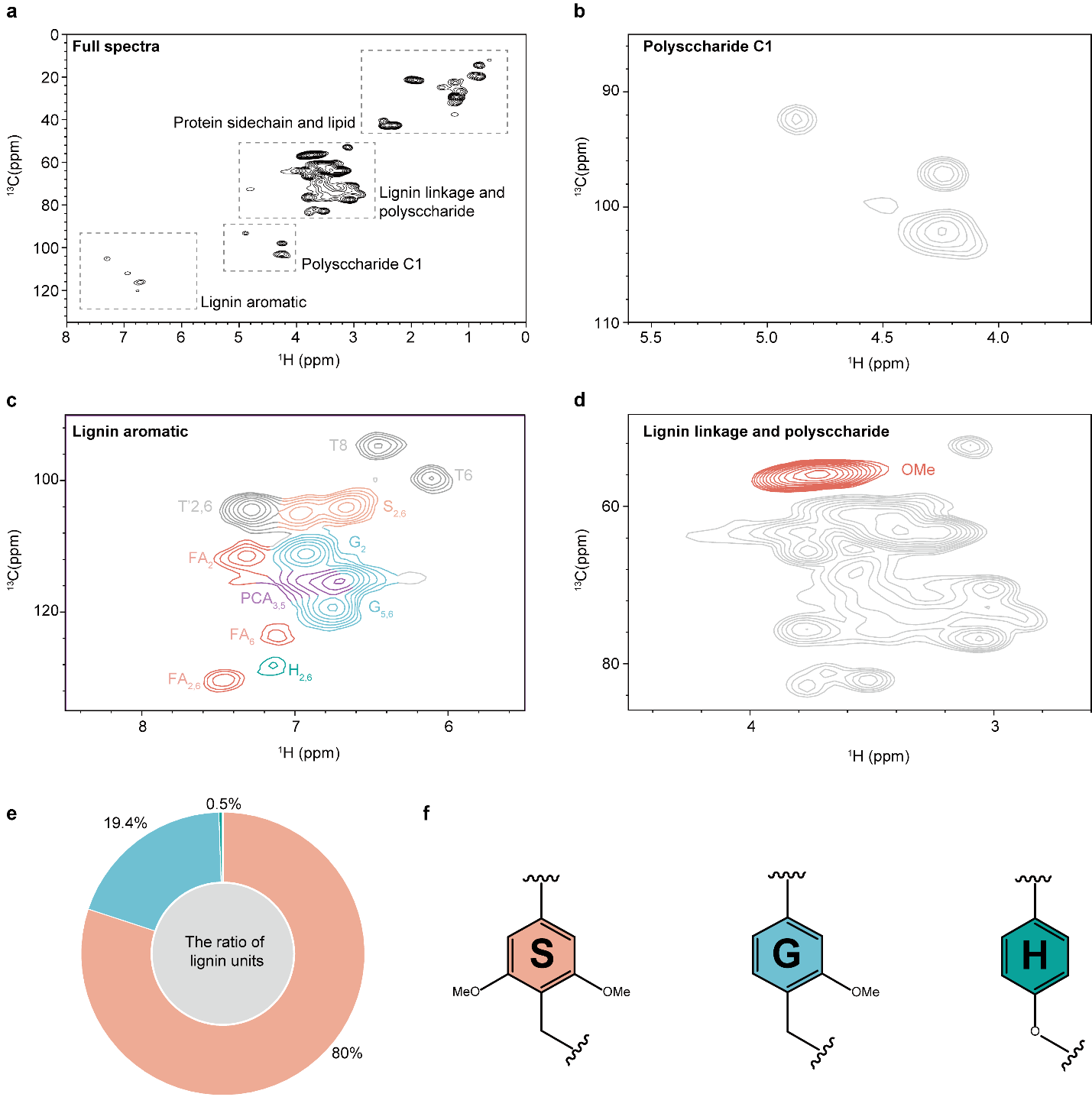

Figure S2. HSQC analysis of lignin and polysaccharides. **a**, A full HSQC spectrum of the wheat straw. The profiles include selected regions: **b**, Focusing on mainly polysaccharide signals; **c**, Highlighting predominantly lignin aromatic peaks; **d**, Featuring primarily lignin linkages and polysaccharide regions. **e**, The ratio of lignin determined from the HSQC; The methods for calculating semi-quantitative values for lignin units are based on procedures outlined in the literature ^1–3^. **f**, Illustrates the structure of lignin units. Main structure ^2^: (S) syringyl units; (G) guaiacyl units; (H) *p*-hydroxyphenyl units; ( *p*-CA) p-coumarates; (FA) ferulates; (T) a likely incorporation of tricin into the lignin polymer through a G-type β-O-4 linkage.

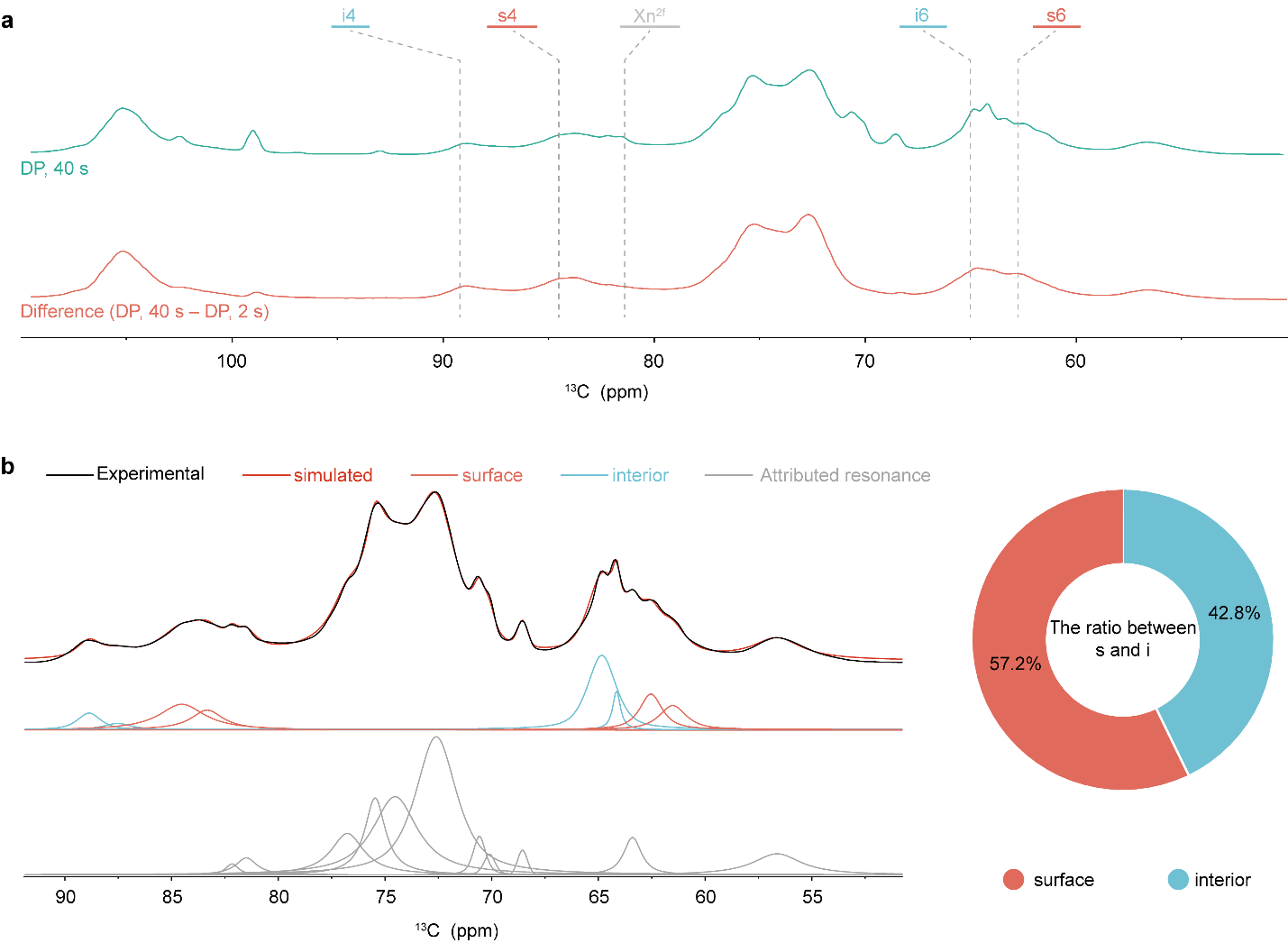

Figure S3. The component analyses of wheat straw using ssNMR. **a**, quantitative analysis is detected by the quantitative DP with 40 s recycle delays, and rigid components can be selectively observed using a difference spectrum. The chemical shifts at C4 and C6 can distinguish the signal. **b**, The surface and interior cellulose region deconvolutions are shown for a DP 40 s spectrum. The ratio of the surface (s) to the sum of the surface (s) and interior (i) can be calculated using integration. (Details see the **Supplementary Table. 4**)­­

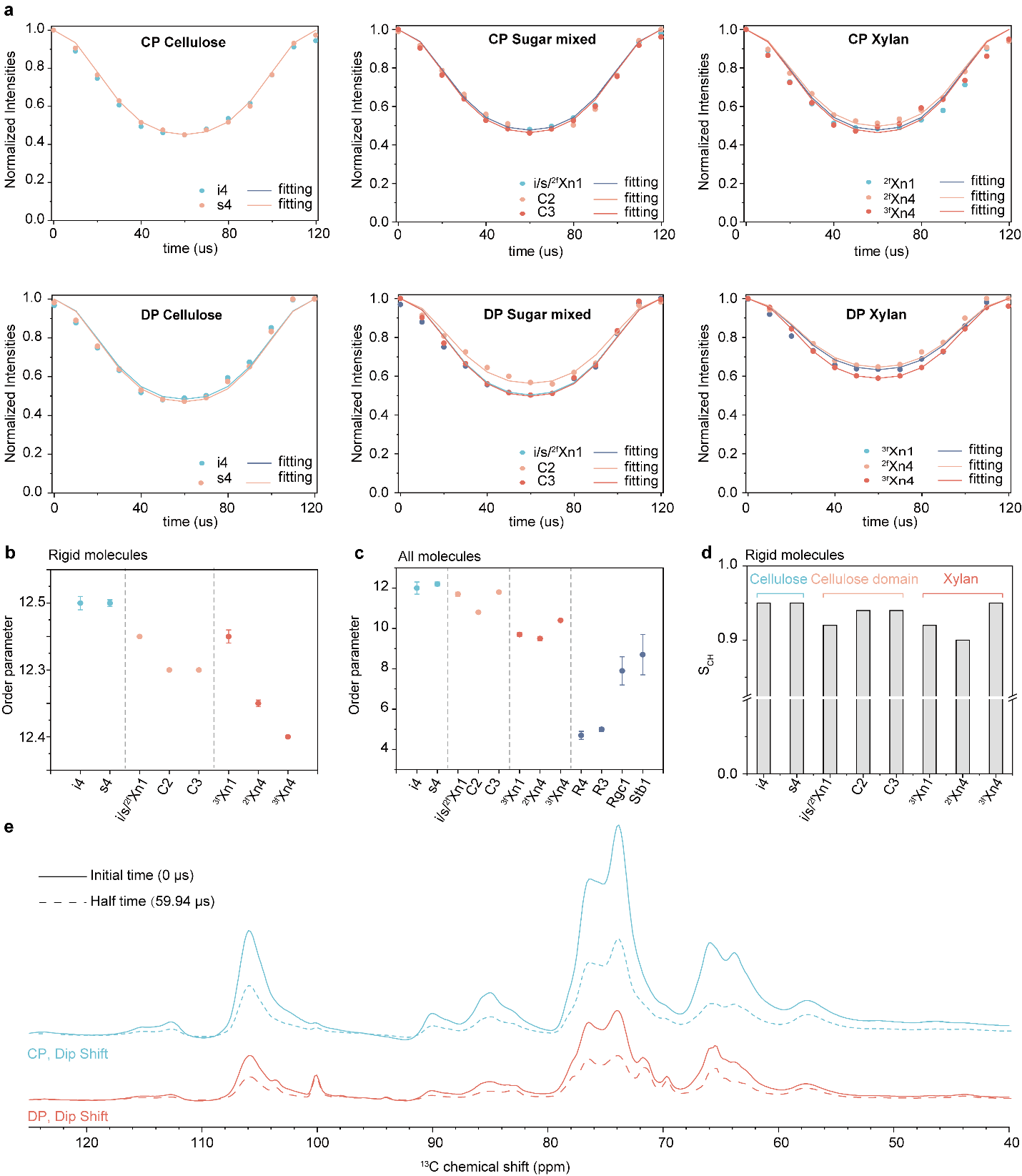

Figure S4. ^13^C-^1^H dipolar order parameters of biopolymers for CP and 40 s DP. **a**, CP-dipolar dephasing spectra (top panel) and 40 s DP-dipolar dephasing spectra (bottom panel) of carbohydrates are shown. The order parameter of rigid molecules **b** and **c**. **d**, The order parameter of S_CH_ was analyzed using CP-DIPSHIFT MAS NMR analysis. **e**, Slices taken at the initial point (0 μs dipolar dephasing) and at half-time (59.94 μs dipolar dephasing) from the 2D DIPSHIFT experiment demonstrate the degree of spectral intensity reduction occurring at half a rotor period.

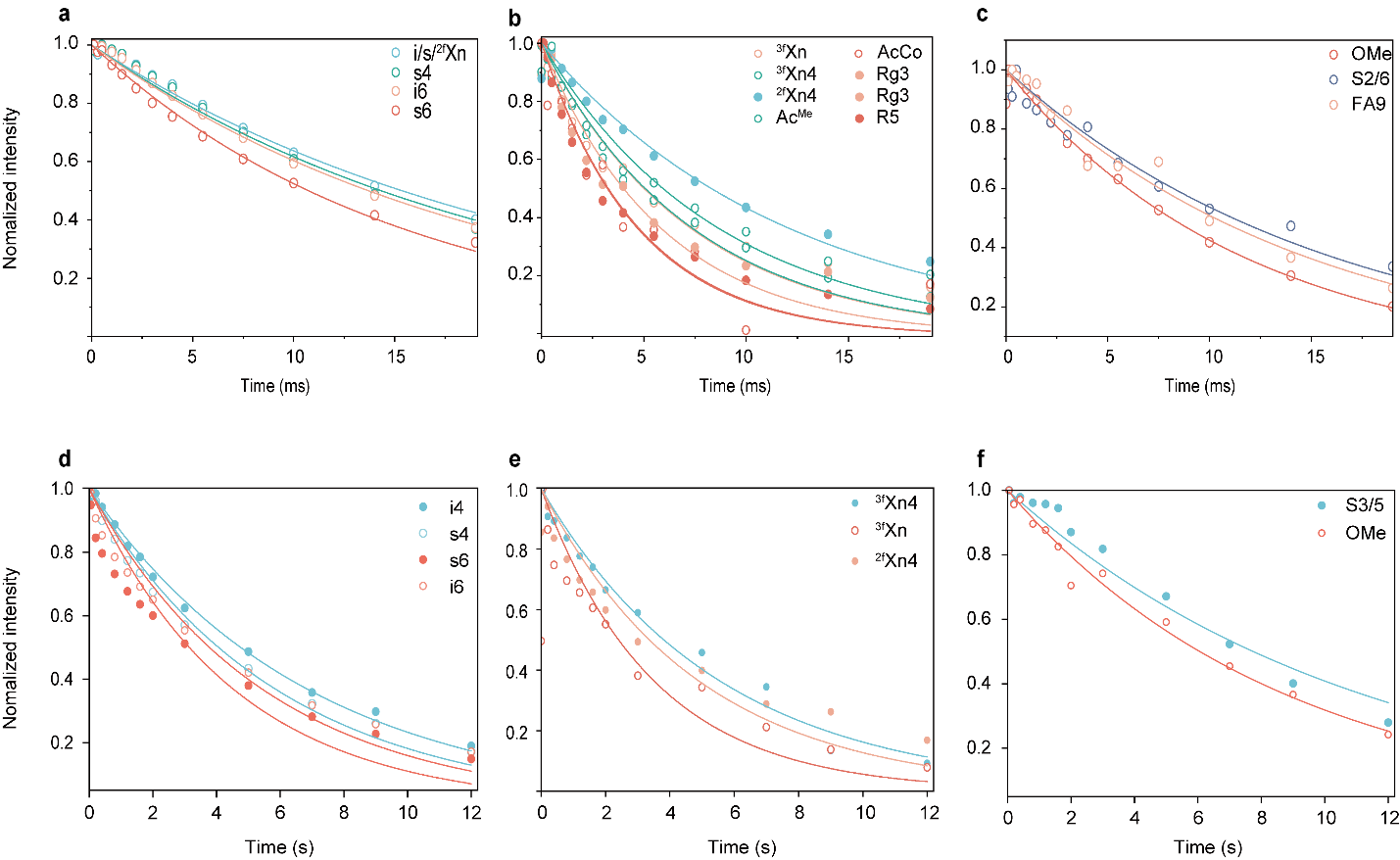

Figure S5. The dynamical behavior of molecules resolved using ssNMR relaxations. This figure elucidates the molecular dynamics by employing ssNMR relaxation techniques. The analysis focuses on three major biopolymers: cellulose, hemicellulose, and lignin, each exhibiting distinct relaxation behaviors indicative of their structural environments and mobility. The ^1^H-T_1ρ_ relaxation curves were analyzed for (**a**) cellulose (interior/surface glucan chains), (**b**) hemicellulose (two/three-fold xylan and other matrices), and (**c**) lignin. These relaxation data were fitted with a single exponential equation. The ^13^C-T1 relaxation curves for (**d**) cellulose (interior/surface glucan chains), (**e**) hemicellulose (two/three-fold xylan), and (**f**) lignin were detected using Torchia CP.

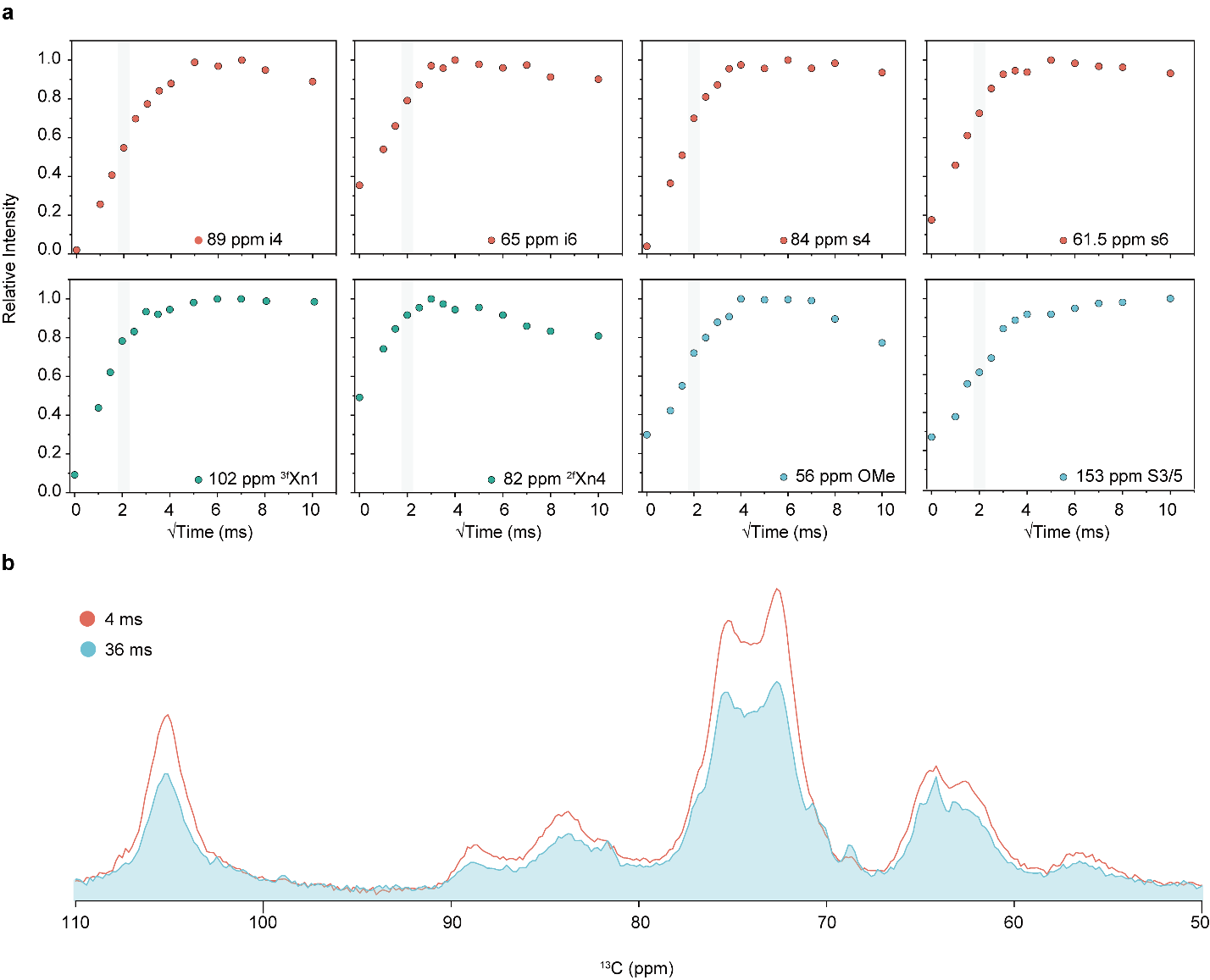

Figure S6. Water to biomolecules buildup spectrums. **a**, The ^1^H spin diffusion curves for water in cellulose, xylan, and lignin are illustrated. The dashed lines indicate the intensities of water-to-carbon transfer observed at 4-ms ^1^H mixing. Since interior cellulose is not present on the microfibril surface, water polarization transfer likely occurs through the surface cellulose via the microfibril’s ^1^H-^1^H dipolar coupling network ^4^. The wheat straw has a fast spin diffusion from water, with ~60–80% of the equilibrium intensity measured at 4-ms ^1^H mixing. **b**, Water-edited spectra are shown, detected at 4-ms and 36-ms mixing times, with the intensities indicating well-hydrated polymers.

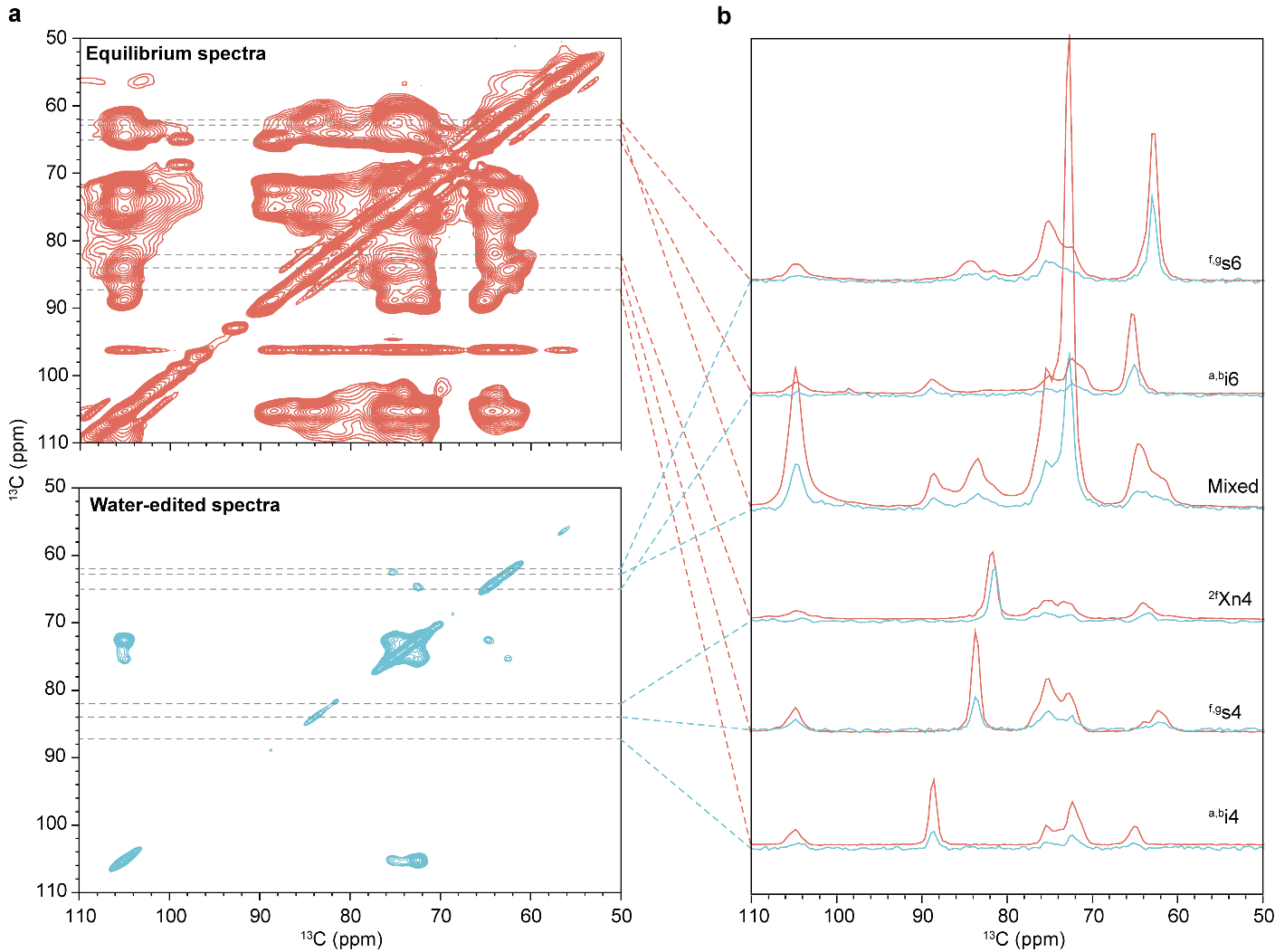

Figure S7. The spectra of accessibility to water. **a**, Water associations are shown in equilibrium spectra (orange) and water-edited spectra (blue). **b**, The representative one-dimensional slices from water-edited (blue) and equilibrium (red) mixing time spectra. The enhanced intensity of the three-fold and two-fold xylan in the water-edited spectra suggests that these structures have stronger interactions with water molecules.

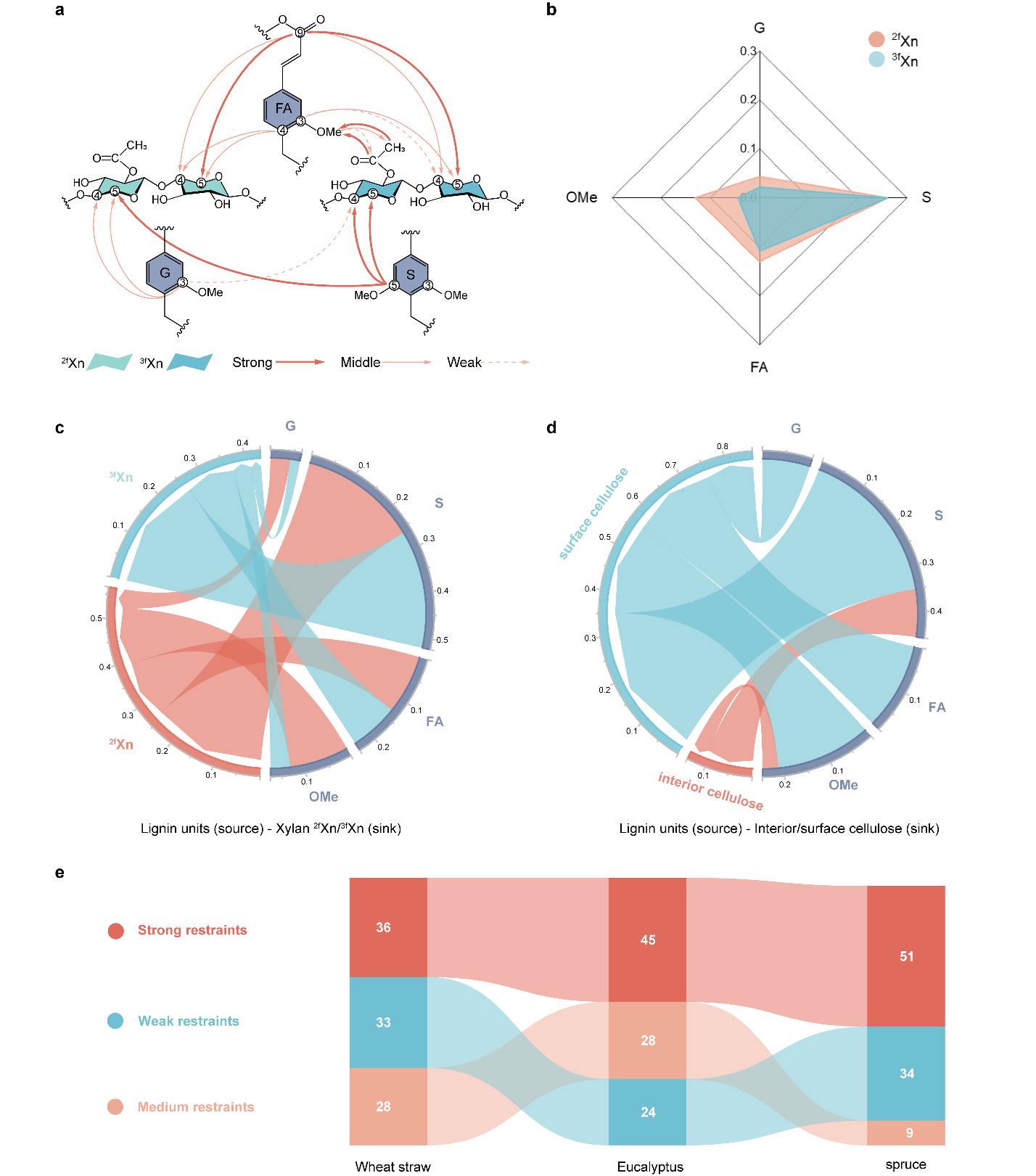

Figure S8. The interaction of polymers in the cell wall. **a**, NMR restraints-observed of lignin-xylan packing was visualized in secondary cell walls; The interaction of lignin-xylan is shown partially (Thick lines represent strong intensity, thin lines medium intensity, and dashed lines weak intensity). **b**, Calculated percentage of intensity from source (Lignin) to sink (^2f^Xn/^3f^Xn) depicted on a polar plot. (**c**) and (**d**) The interactions of polysaccharide-lignin in the wheat straw were plotted; Each arc represents a source or sink, while the lines connecting the circles illustrate the interactions between the two components. **e**, The intensities of intermolecular cross-peaks from different lignocellulosic biomass ^5^. The wheat straw shows relatively weak restrain compared with other woody biomass materials (eucalyptus (*Eucalyptus grandis*) and spruce (*Picea abies*)).

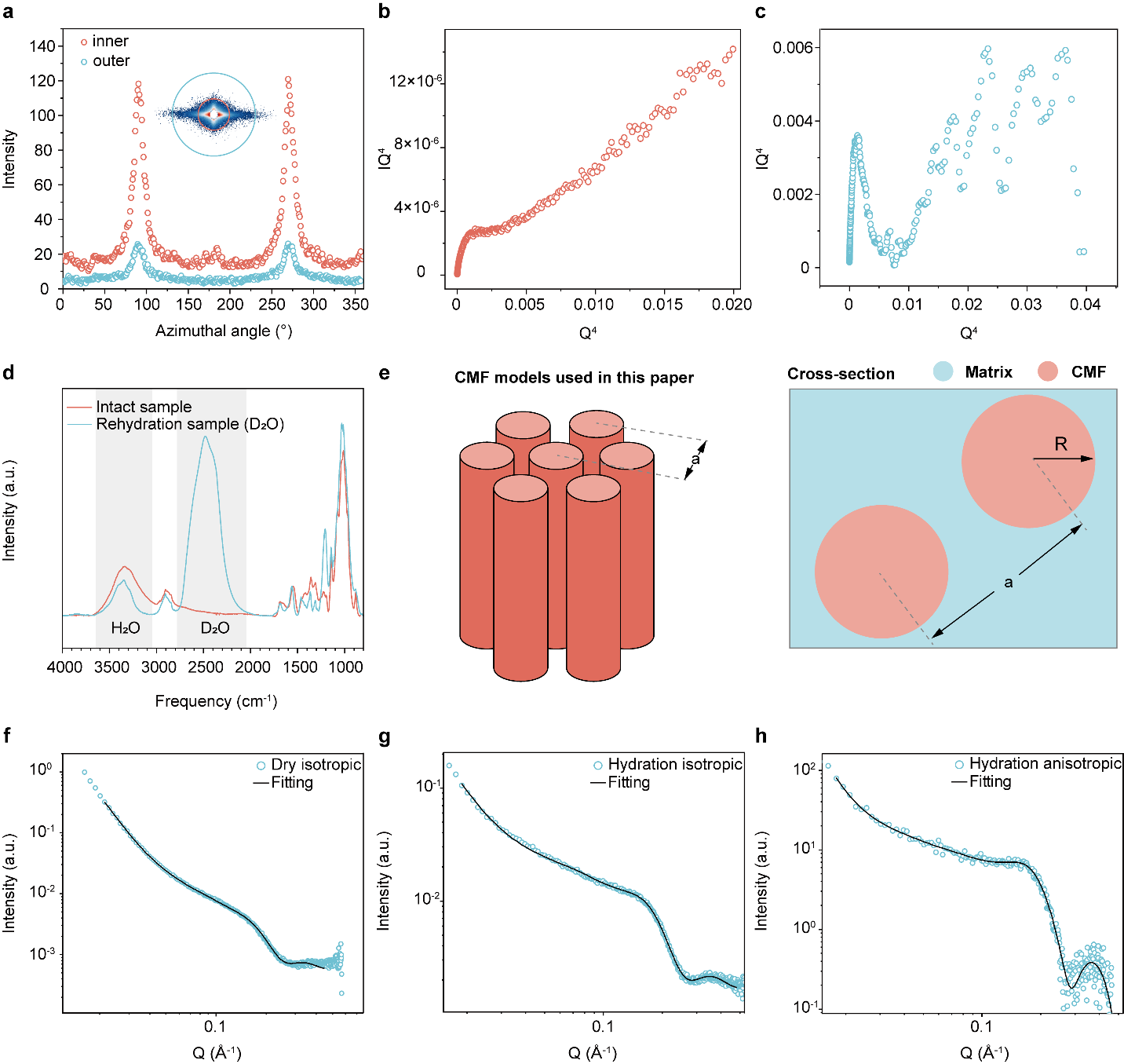

Figure S9. The model fitting and architecture of the fibrils. **a**, Azimuthal intensity profiles at different q values. (**b**) and (**c**) Porod analysis (Q^4^ × I (Q) vs. Q^4^) for isotropic and anisotropic samples of intact was conducted. a linear was observed at the values of q greater than 0.23 Å^-1^, corresponding to a particle size of approximately 27.3 nm in real space. This linearity typically indicates a two-phase system, characterized by the separation between the fibril and the surrounding matrix ^6^. However, in the anisotropic sample, the straight-line behavior disappeared, likely due to the effects of background subtraction. (**d**) FTIR-ATR spectra of the sample in a dry state and after equilibration with D_2_O. These results are by the core-shell model ^7,8^. **e**, The fibril model used in this paper ^9^. In small-angle scattering (SAS), the measured intensity, I(q), can generally be described as the product of a form factor, P(q), and the structure factor, S(q). The form factor P(q) arises from the shape and size of the fibrils within each particle, whereas the structure factor S(q) captures the effects of volume fraction and interparticle distances in systems with ordered arrangements. The P(q) features a characteristic shoulder with a gradually decreasing intensity. In contrast, the S(q) displays a distinct “structure peak,” whose position indicates interparticle spacing. When this structure peak appears in the SAS profile, both form factor and structure factor contributions can be included in the model-fitting process. **f-g**, Fitting curves and data from wheat straw under dry and wet conditions.

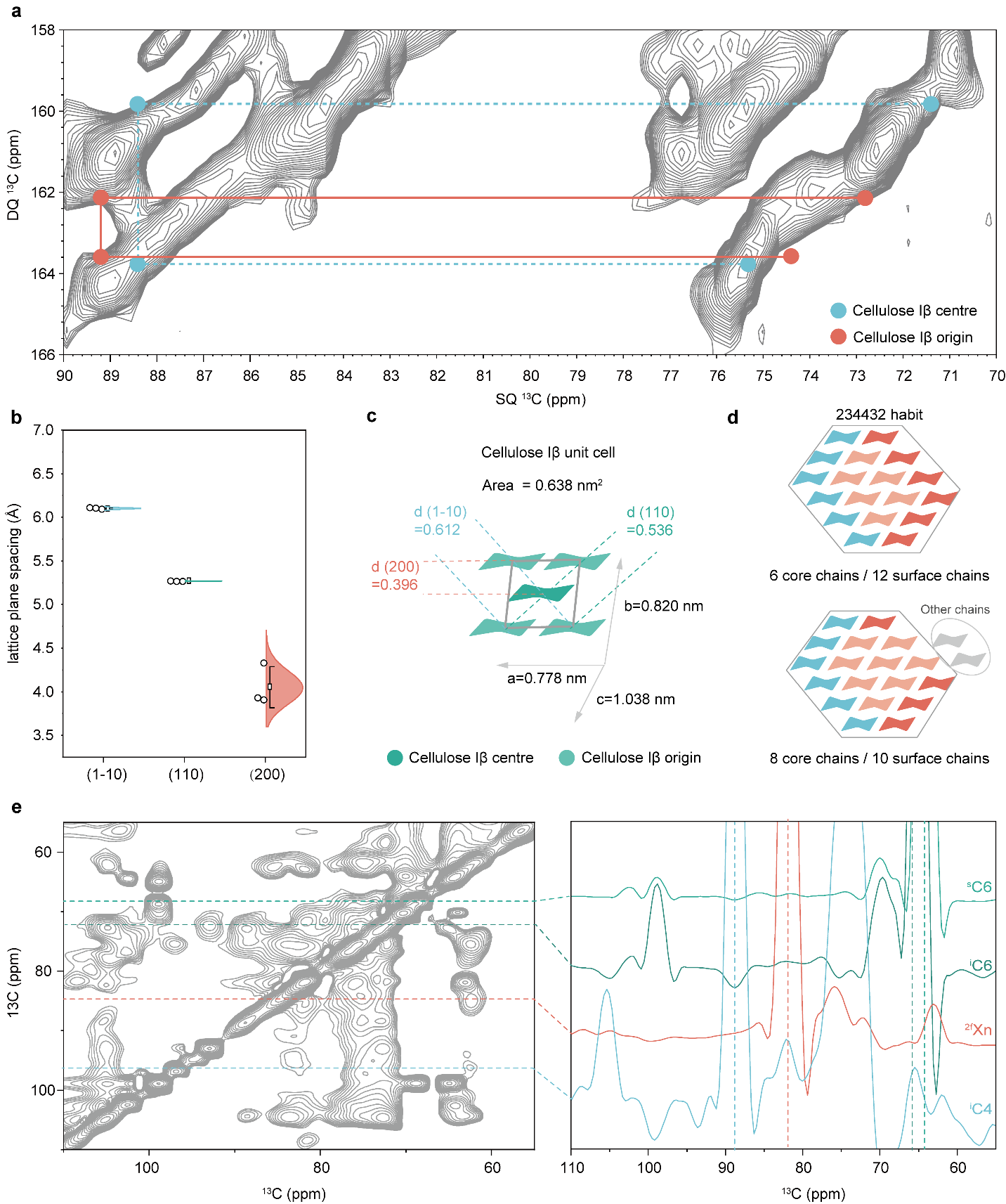

Figure S10. Structural analysis of cellulose fibrils. This figure provides an in-depth structural analysis of cellulose fibrils within the cell wall, utilizing a combination of X-ray diffraction and solid-state Nuclear Magnetic Resonance techniques. The study elucidates cellulose’s crystalline structure, lattice spacing, and molecular interactions. **a**, The cell wall contains cellulose with an Iβ component consistent with the prior work ^10^. **b**, Lattice plane spacing (**d**) was determined by fitting parameters for different planes (200, 110, and 1-10). For each plane, measurements were taken **n = 3** times, and the results were reported as mean values with standard deviations. **c**, Schematic of the monoclinic unit cell of cellulose Iβ. **d**, To maintain structural integrity, ECF accommodates a potential cellulose habit (234432), which requires the incorporation of additional chains as the core chain extends to 8 chains. **e**, Left: The CP-gated PDSD spectrum of wheat straw obtained with a 100 ms mixing time using solid-state NMR (ssNMR). Right: 1D slices extracted from the 2D pattern for further analysis.

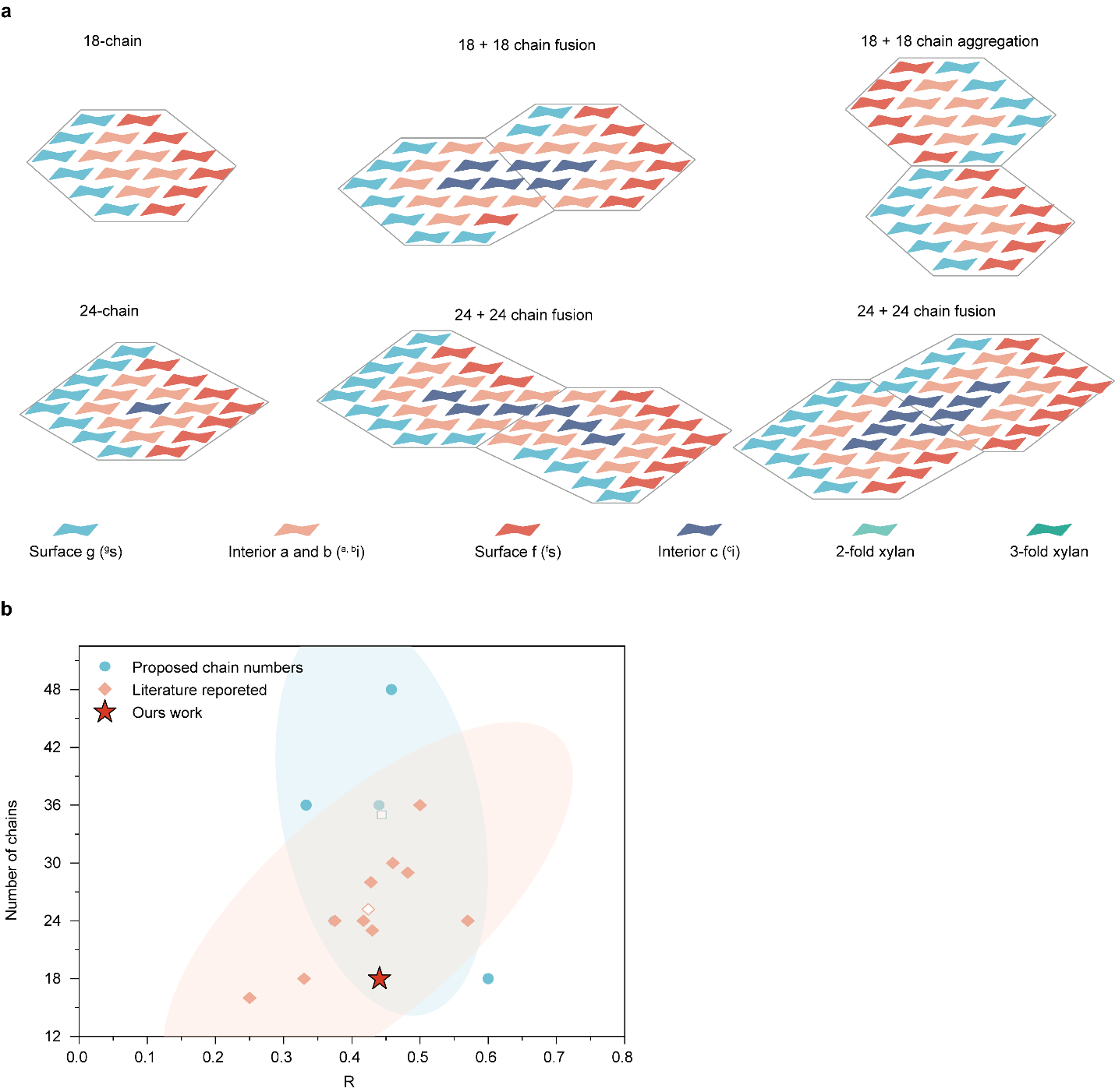

Figure S11. Potential Fibril Models Inferred from ssNMR Insights. **a**, Proposed ECF structural models were established. The ECF model includes 18 and 24-chain horizontal cross-sections in fibril bounding (fused and aggregated states), providing a range of models with different habits. Labels for the hydrophobic surface chains are ^g^s, for the hydrophilic surface chains ^f^s, for the middle layer internal chains ^a,b^i, and for the deeply embedded core chains ^c^i ^11^. **b**, The number of glucan chains in cellulose microfibrils varies according to the interior-to-all glucan chain ratio (R) ^5,12–15^. Data analysis indicates CMFs might consist of diverse chains among various plants. Our reasonable model conforms to its size and shape of measuring, compared with other models fitted by the ratio.

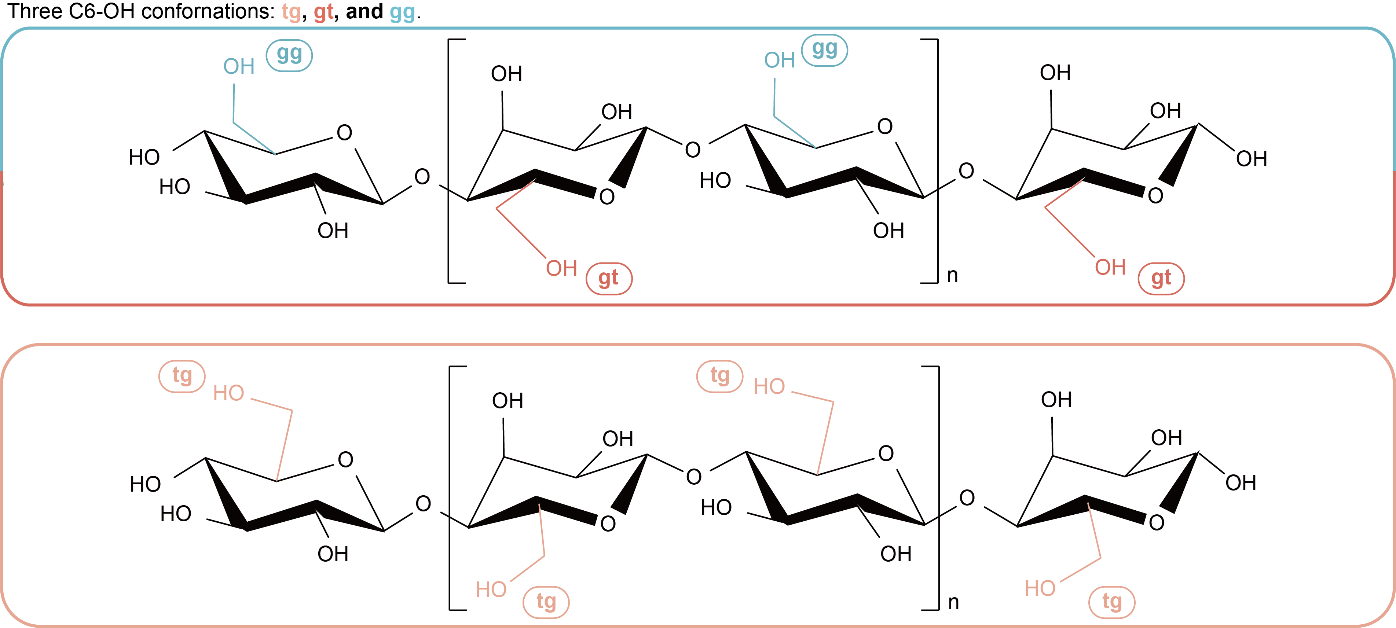

Figure S12. Schematic representation of C6-OH conformations in different cellulose residues.

### Table

Table S1. ^13^C chemical shifts of rigid molecules. All components are identified from ^13^C CP-based INADEQUATE spectra. Superscripts are used to denote different allomorphs. Not applicable (/). Unidentified (-).

| **CP-INADEQUATE** | **Type** | **C1** | **C2** | **C3** | **C4** | **C5** | **C6** |  |  |  | **Ref.** |
| --- | --- | --- | --- | --- | --- | --- | --- | --- | --- | --- | --- |
| Cellulose | ^a,b^i | 105.1 | 72.6 | 75.4 | 88.6 | 72.4 | 64.7 | / | / | / | ^5,16^ |
|  |  | 105.3 | 72.4 | 75.4 | 88.6 | 72.4 | 64.7 | / | / | / |  |
|  | ^c^i | 104.8 | 72.3 | 75.2 | 88.0 | 70.9 | 65.7 | / | / | / |  |
|  | ^f^s | 105.4 | 72.0 | 76.9 | 84.4 | 75.3 | 62.6 | / | / | / |  |
|  | ^g^s | 105.8 | 72.4 | 76.4 | 83.7 | 73.6 | 61.2 | / | / | / |  |
|  | **Type** | **C1** | **C2** | **C3** | **C4** | **C5** | **C6** | **AC^CO^** | **AC^Me^** |  | **Ref.** |
| Xylan | ^2f,a^ Xn | 105.2 | 72.4 | 73.0 | 82.0 | 63.9 | / | 173.7 | 20.4 | / | ^17^ |
|  | ^2f,b^ Xn | 105.1 | 72.3 | 75.7 | 81.0 | 63.6 | / |  |  | / |  |
|  | ^2f,c^ Xn | 105.0 | 72.5 | 75.3 | 80.0 | - | / |  |  | / |  |
|  | ^2f,d^ Xn | 105.1 | 72.5 | 76.2 | 78.9 | - | / |  |  | / |  |
|  | Xn | 105.0 | 72.4 | 74.2 | 78.2 | - | / |  |  | / |  |
|  | ^3f,a^ Xn | 102.5 | 72.2 | 74.2 | 77.7 | 63.2 | / |  |  | / |  |
|  | ^3f,b^ Xn | 102.0 | 72.8 | 73.6 | 77.4 | 62.9 | / |  |  | / |  |
|  | **Type** | **C1** | **C2** | **C3** | **C4** | **C5** | **C6** | **C7** | **C8** | **C9** | **Ref.** |
| Lignin | S_a_ | 134.0 | 103.9 | 154.1 | 134.9 | 154.1 | 103.9 | / | / | / | ^5^ |
|  | S_b_ | - | 106.7 | 154.3 | 137.5 | 154.3 | 106.7 | / | / | / |  |
|  | G | - | 109.5 | 151.9 | 147.9 | 115.6 | - | / | / | / |  |
|  | FA | - | - | 151.9 | 151.9 | 113.7 | - | - | 113.7 | 169.9 |  |
|  | H | 128.9 | 126.0 | 121.7 | 163.9 | 121.7 | 126.0 | / | / | / |  |

Note: The assignment of chemicals can also refer to the Complex Carbohydrates Magnetic Resonance Database (CCMRD) ^18^.

Table S2. Semi-quantitation of the molar composition of cellulose and xylan. Cellulose in these different chemical environments contains interior glucan chains (^a,b^i: middle layer; ^c^i: embedded core chains) and surface glucan chains (^f^s: hydrophilic surface; gs: hydrophobic surface). Xylan in these different environments contains xylose chains (^2f^Xn: two-fold xylan; ^3f^Xn: three-fold xylan; Xn: mixed conformation).

| **Cellulose** | | | | | |
| --- | --- | --- | --- | --- | --- |
| interior cellulose (i) % | | surface cellulose (s) % | | | R |
| ^a,b^i | ^c^i | ^f^s | | ^g^s | i/(i+s) % |
| 43.5 | 13.2 | 32.5 | | 10.8 | 56.7 |
| **Xylan** | | | | | |
| ^2f^Xn % | | | Xn % | ^3f^Xn % | |
| 73.2 | | | 13.0 | 13.9 | |

Note: The calculation formula is as follows ^5,21^:

^a,b^i% = ^a,b^i/(^a,b^i + ^c^i + ^f^s+ ^g^s) × 100%;

^c^i% = ^c^i/(^a,b^i + ^c^i + ^f^s + ^g^s) × 100%;

^f^s% = ^f^s/(^a,b^i + ^c^i + ^f^s + ^g^s) × 100%;

^g^s% = ^g^s / (^a,b^i + ^c^i + ^f^s + ^g^s) × 100%;

^2f^Xn% = ^2f^Xn/(^2f^Xn + Xn + ^3f^Xn) × 100%;

Xn% = Xn/(^2f^Xn + Xn + ^3f^Xn) × 100%;

^3f^Xn% = ^3f^Xn/(^2f^Xn + Xn + ^3f^Xn) × 100%.

The areas of the well-resolved peak pairs, which are distinguishable in the CP J-INADEQUATE spectra, were selected for integration because these peaks are distinct and do not exhibit any overlay:

^a,b^i: the average of ^a,b^i5 and ^a,b^i6;

^c^i: the average of ^c^i5 and ^c^i6;

^f^s: the average of ^f^s5 and ^f^s6;

^g^s: the average of ^g^s5 and ^g^s6;

^2f^Xn: the average of ^2f^Xn4 and ^2f^Xn5;

Xn: the average of Xn4 and Xn5;

^3f^Xn: the average of ^3f^Xn4 and ^3f^Xn5.

Table S3. ^13^C chemical shifts of mobile molecules. All components are identified from ^13^C DP-based INADEQUATE spectra. Superscripts are used to denote different allomorphs. Weak signals or minor species are indicated using “w” Not applicable (/). Unidentified (-).

| **DP-INADEQUATE** | **Type** | **C1** | **C2** | **C3** | **C4** | **C5** | **C6** | **OMe** | **Ref.** |
| --- | --- | --- | --- | --- | --- | --- | --- | --- | --- |
| Arabinose | A | 105.2 | 82.8 | 76.5 | 82.1 | 62.1 | / | / |  |
| Glucuronic acid | Glu | 98.0 | 74.9 | - | - | - | - | / |  |
| Starch | St^a^ | 92.7 | 72.0 | 75.3 | 76.3 | 70.5 | 62.5 | / | ^19^ |
|  | St^b^ | 96.6 | 74.6 | 77.4 | 63.4 | 72.12 | 61.6 | / |  |
| Rhamnose units | R | 102.4 | 76.7 | 72.0 | 70.4 | 68.4 | - | / | ^19,20^ |
|  | R | 104.3 | 77.5 | 72.0 | 70.4 | 68.4 | - | / |  |
| Galactose units_ambiguous |  | 99.0 | 64.4 | 82.9 | 76.65 | 74.88 | 62.1 | / | ^19^ |
| Xylan | ^2f,a^Xn | 105.6 | 71.6 | 75.0 | 81.29 | 62.9 | / | / |  |
| Lignin | S_a_ | 140.44 (w) | 103.9 | 154.1 | 134.9 | 154.1 | 103.9 | 56.1 |  |
|  | S_b_ | - | 106.7 | 154.3 | 137.5 | 154.3 | 106.7 | 56.1 |  |
|  | G | - | - | 149.6 | 149.6 | - |  | - |  |
|  | FA | - | - | 149.6 | 149.6 | 113.71 |  | - |  |

Table S4. Deconvolution details of 1D 40s quantitative 13C ssNMR spectra in the polysaccharide region.

| **Carbon site (ppm)** | **Height** | **Integral area** |
| --- | --- | --- |
| 64.1 | 4391918.0 | 33108059.4 |
| 68.6 | 2768728.1 | 18742487.1 |
| 70.1 | 2317466.6 | 21463438.1 |
| 70.6 | 4315875.7 | 45636902.6 |
| 82.2 | 1118976.9 | 15083047.3 |
| 63.4 | 4183356.7 | 71600552.4 |
| 75.5 | 8582933.0 | 179840373 |
| 87.5 | 673357.9 | 13195099.8 |
| 81.5 | 1793615.2 | 40122345.9 |
| 62.5 | 3995521.6 | 85787537.5 |
| 88.9 | 1848251.5 | 40023004.6 |
| 61.5 | 2679947. | 68420624.1 |
| 64.8 | 8259910.1 | 202691935.4 |
| 83.3 | 2176490.1 | 71970449.51 |
| 76.8 | 4631286.1 | 157063459.8 |
| 72.6 | 15390293.5 | 591355090.5 |
| 74.5 | 8715237.8 | 389543184.3 |
| 84.5 | 2834880.0 | 133818966.8 |
| 56.6 | 2359475.3 | 117881981.9 |

Table S5. The dipolar order parameter S_CH_.^13^C-^1^H dipolar coupling and dipolar order parameters of biopolymers for ^13^C CP and quantitative DP. Unidentified (-). Error bars are standard errors calculated from the ratio of signal/noise.

| **Site** | **Dip Coupling,**  **CP** | **S_CH_,**  **CP** | **Dip Coupling,**  **DP 40s** | **S_CH_,**  **DP 40s** |
| --- | --- | --- | --- | --- |
| i4 | 12.5±0.2 | 0.95 | 12.0±0.3 | 0.92 |
| s4 | 12.5±0.1 | 0.95 | 12.2±0.1 | 0.93 |
| i/s/^2f^ Xn | 12.1±0 | 0.92 | 11.7±0.1 | 0.89 |
| C2 | 12.3±0 | 0.94 | 10.8±0 | 0.82 |
| C3 | 12.3±0 | 0.94 | 11.8±0 | 0.9 |
| ^3f^Xn1 | 12.1±0.2 | 0.92 | 9.7±0.1 | 0.74 |
| ^2f^Xn4 | 11.8±0.1 | 0.90 | 9.5±0.1 | 0.73 |
| ^3f^Xn4 | 12.4±0 | 0.95 | 10.4±0 | 0.79 |
| R4 | - | - | 4.7±0.2 | 0.33 |
| R3 | - | - | 5.0±0.1 | 0.35 |
| Rgc1 | - | - | 7.9±0.7 | 0.55 |
| Stb1 | - | - | 8.7±1.0 | 0.61 |

Table S6. ^1^H-T1ρ relaxation times of lignin and polysaccharides. The data are fit using a single exponential equation: 𝐼 (𝑡) = 𝑒^−𝑡/𝑇^. Standard deviations of the fitting parameters are used as error bars.

| **Site** | **Atom (ppm)** | **T1, CP (ms)** |
| --- | --- | --- |
| FA9 | 169.9 | 14.8±0.8 |
| G/FA 3/4 | 149.6 | 7.9±1.9 |
| S2/6 | 103.9 | 16.1±0.8 |
| OMe | 56.3 | 11.7±0.2 |
| i/s/^2f^Xn1 | 105 | 22.2±0.7 |
| i4 | 88.6 | 31.8±2.9 |
| s4 | 83.7 | 20.7±0.7 |
| i6 | 64.6 | 19.8±0.4 |
| s6 | 62.4 | 15.5±0.3 |
| AcCo | 173.5 | 4.8±0.5 |
| Ara (a) 1 | 107.8 | 13.9±0.8 |
| Ara (b) 1 | 109.7 | 14.4±1.1 |
| ^3f^Xn1 | 102.5 | 7.3±0.6 |
| ^2f^Xn4 | 81.8 | 12.1±0.4 |
| ^3f^Xn4 | 77.7 | 8.7±0.5 |
| ^2f,3f^Xn5 | 63 | 3±0.3 |
| AcMe | 20.9 | 7.4±0.4 |
| Rgc1 | 98.6 | 1.4±0.1 |
| Rg3 | 69.9 | 5.9±0.4 |
| R5 | 68.7 | 4.7±0.3 |

Table S7. ^13^C­-T1 relaxation times of lignin and polysaccharides. The data are fit using single exponential equations 𝐼 (𝑡) = 1−2𝑒^−𝑡/𝑇^ for T1, DP (inversion recovery), and 𝐼 (𝑡) = 𝑒^−𝑡/𝑇^ for T1, CP (Torchia CP). Standard deviations of the fitting parameters are used as error bars. Unidentified (-).

| **Type** | **Atom (ppm)** | **T1, CP (s)** | **Atom (ppm)** | **T1, DP (s)** |
| --- | --- | --- | --- | --- |
| FA9 | 169.9 | 3.9±0.6 | 169.9 | 5±0.4 |
| S3/5 | 153 | 8.2±0.7 | 154.5 | 4.9±0.5 |
| G/FA 3/4 | - | - | 148.5 | 4.9±0.5 |
| S4 | - | - | 135 | 6.4±0.7 |
| S2/6 | 103.9 | 4.0±0.5 | 103.8 | 6±0.5 |
| OMe | 56.1 | 8.7±0.3 | 56.4 | 4.6±0.5 |
| i/s/^2f^Xn1 | 105 | 5.8±0.3 | 105 | 6.2±0.5 |
| i4 | 88.6 | 6.9±0.1 | 88.6 | 7.1±0.6 |
| s4 | 84 | 5.9±0.1 | 83.6 | 5.4±0.5 |
| i6 | 65 | 5.5±0.4 | 65 | 3.3±0.5 |
| s6 | 62.5 | 4.5±0.4 | 62.7 | 3.2±0.5 |
| AcMe | - | - | 21 | 5.2±0.5 |
| AcCo | - | - | 173 | 4.9±0.5 |
| Ara (a) 1 | - | - | 107.9 | 5.2±0.6 |
| Ara (b) 1 | - | - | 109.7 | 5.4±0.8 |
| ^3f^Xn1 | 102.5 | 3.5±0.3 | 102.4 | 2.3±0.3 |
| ^2f^Xn4 | 82 | 4.9±0.4 | 82.6 | 3.6±0.5 |
| ^3f^Xn4 | 77.7 | 5.5±0.3 | 77 | 3.3±0.4 |
| ^2f^Xn5 | 64 | 4.4±0.4 | 64 | 1.6±0.3 |
| ^3f^Xn5 | 63 | 4±0.4 | 63 | 2.5±0.4 |
| Rgc1 | - | - | 98.7 | 0.9±0.1 |
| Sta1 | - | - | 92.8 | 0.5±0.1 |
| Stb1 | - | - | 96.7 | 0.5±0.1 |
| Rg3 | - | - | 70 | 0.8±0.1 |
| R5 | - | - | 68.4 | 0.8±0.1 |

Table S8. Water-edited intensities of biopolymers. The intensity ratio is derived by comparing water-edited and control 2D correlation spectra. Error bars represent the standard deviations of the NMR signal-to-noise ratios.

| **Type** | **Carbon site** | **Raletive Intensity (S/S0)** | **Average value** |
| --- | --- | --- | --- |
| Interior cellulose | i4-1 | 0.27±0.03 | 0.33 |
|  | i4-3 | 0.32±0.01 |  |
|  | i4-2/5 | 0.30±0.01 |  |
|  | i6-1 | 0.36±0.01 |  |
|  | i6-3 | 0.38±0.00 |  |
|  | i6-2/5 | 0.32±0.01 |  |
| Surface cellulose | s4-1 | 0.42±0.01 | 0.36 |
|  | s4-3 | 0.36±0.03 |  |
|  | s4-5 | 0.30±0.05 |  |
|  | s4-2 | 0.38±0.03 |  |
|  | s4-6 | 0.34±0.04 |  |
| two-fold xylan | Xn^2f^4-3 | 0.35±0.04 | 0.36 |
|  | Xn^2f^4-2 | 0.43±0.01 |  |
|  | Xn^2f^4-5 | 0.65±0.01 |  |
|  | Xn^2f^4-1 | 0.42±0.02 |  |
|  | Xn^2f^3-5 | 0.43±0.01 |  |
|  | Xn^2f^5-1 | 0.22±0.1 |  |
|  | Xn^2f^5-4 | 0.27±0.08 |  |
| Mixed | Xn^3f,2f^2/3-1 | 0.31±0.07 | 0.33 |
|  | Xn^3f,2f^2/3-4 | 0.36±0.04 |  |
|  | Xn^3f,2f^2/3-5 | 0.33±0.06 |  |
| three-fold xylan | Xn^3f^5-1 | 0.53±0.00 | 0.4 |
|  | Xn^3f^5-4 | 0.29±0.07 |  |
|  | Xn^3f^4-1 | 0.42±0.02 |  |
|  | Xn^3f^4-2 | 0.34±0.06 |  |
|  | Xn^3f^4-3 | 0.45±0.02 |  |
|  | Xn^3f^4-5 | 0.37±0.04 |  |
| Lignin-OMe | OMe-S1/4a | 0.40±0.00 | 0.31 |
|  | OMe-S3/5 | 0.34±0.02 |  |
|  | OMe-G3 | 0.25±0.06 |  |
|  | OMe-G4 | 0.23±0.07 |  |

Table S9. Intermolecular interactions of polymers. In total, 97 restraints are identified, including 36 strong restraints, 28 medium ones, and 33 weak restraints.

| **Interaction type** | **Strong restraints** | **Medium restraints** | **Weak restraints** | **Total** |
| --- | --- | --- | --- | --- |
| Cellulose-Xylan Ac | 1 | 0 | 2 | 3 (3.1%) |
| Lignin-Cellulose | 4 | 3 | 9 | 16 (16.5%) |
| Lignin-Lignin | 8 | 11 | 6 | 25 (25.5%) |
| Lignin-Xylan | 11 | 13 | 15 | 39 (40.2%) |
| Lignin-Mixed sugar | 12 | 1 | 1 | 14 (14.4%) |
| Sum | 36 | 28 | 33 | 97 (100%) |
| **Percentage of the interaction site** | | | | |
| 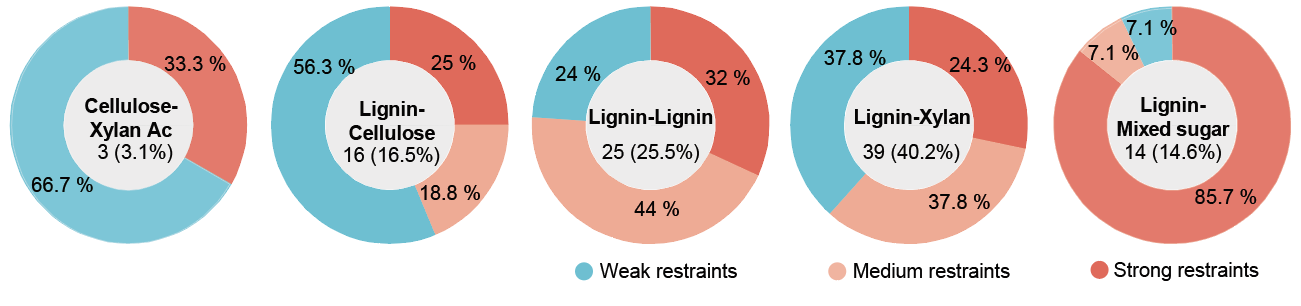 | | | | |

### Table S10. The fitting parameters of fibril in the cell wall.

| **Fitting parameters** | | **Anisotropic** | | **Isotropic** | |
| --- | --- | --- | --- | --- | --- |
|  |  | **Dry** | **Hydration** | **Dry** | **Hydration** |
| Radius (Å) | | 12.9±0.6 | 13.2±0.4 | 13.6±1.8 | 13.3±0.7 |
| ΔR/‾R | | 0.1±0.1 | 0.2±0.1 | 0.1±0.0 | 0.2±0.1 |
| a (Å) | | 28.6±2.9 | 27.0±5 | 31.0±1.1 | 29.8±2.3 |
| Δa/‾a | | 0.5±0.1 | 0.5±0.2 | 0.3±0.0 | 0.4±0.1 |
| Rc | | 9.1±0.1 | 9.4±0.0 | 9.6±0.1 | 9.4±0.2 |
| Cross-section area for cellulose microfibril core (Minimum, nm^2^) | | 2.0±0.1 | 2.1±0.0 | 2.3±0.1 | 2.1±0.1 |
| Cross-section area for cellulose microfibril core (Maximum, nm^2^) | | 2.5±0.1 | 2.7±0.0 | 2.9±0.1 | 2.7±0.1 |
| Core chain number N_core_ (Minimum) | | 6.2±0.3 | 6.6±0.1 | 7.2±0.4 | 6.7±0.3 |
| Core chain number N_core_ (Maximum) | | 7.8±0.3 | 8.3±0.1 | 9.0±0.4 | 8.4±0.4 |
| Surface chain number N_shell_ (minimum) | | 10.2±0.3 | 9.6±0.2 | 9.0±0.4 | 9.6±0.4 |
| Surface chain number N_shell_ (Maximum) | | 11.8±0.3 | 11.4±0.1 | 10.8±0.4 | 11.3±0.3 |
| The estimated R values | 18 chain number | 34.4%–43.3% | 36.9%–46.4% | 40%–50% | 37.2%–46.8% |
|  | 24 chain number | 25.8%–32.5% | 27.7%–34.5% | 30%–37.5% | 27.9%–35.2% |
|  | 36 chain number | 17.2%–21.7% | 18.4%–23.1% | 20%–25% | 18.6%–23.4% |

Note: N= N_core_/R; The estimated R values, R = N_Core_/(chains number). For all models, n= 3. Values (Radius, ΔR/‾R, a, and Δa/‾a) are presented as mean±RMSE (Root Mean Square Error), while other values are given as mean±s.e.

Table S11. The crystalline structure of fibril (n=3). Values are presented as mean±s.e.

| **Crystalline plane** | **1–10** | **110** | **200** |
| --- | --- | --- | --- |
| Crystalline structure (L, nm) | 2.4±0.2 | 2.6±0.0 | 2.7±0.2 |
| Lattice plane spacing (d, Å) | 6.1±0.0 | 5.3±0.0 | 4.3±0.2 |
